# Metabolic selection of a homologous recombination mediated loss of glycosomal fumarate reductase in *Trypanosoma brucei*

**DOI:** 10.1101/2020.11.29.403048

**Authors:** Marion Wargnies, Nicolas Plazolles, Robin Schenk, Oriana Villafraz, Jean-William Dupuy, Marc Biran, Sabine Bachmaier, Hélène Baudouin, Christine Clayton, Michael Boshart, Frédéric Bringaud

**Affiliations:** Univ. Bordeaux, CNRS, Microbiologie Fondamentale et Pathogénicité (MFP), UMR 5234, F-33000 Bordeaux, France; Univ. Bordeaux, CNRS, Centre de Résonance Magnétique des Systèmes Biologiques (CRMSB), UMR 5536, F-33000 Bordeaux, France; Fakultät für Biologie, Genetik, Ludwig-Maximilians-Universität München, Großhaderner Strasse 2-4, D-82152 Martinsried, Germany; Univ. Bordeaux, Plateforme Protéome, F-33000, Bordeaux, France; Zentrum für Molekulare Biologie der Universität Heidelberg (ZBMH), Universität Heidelberg, Im Neuenheimer Feld 282, 69120 Heidelberg, Germany

**Keywords:** *Trypanosoma*, genomic rearrangement, homologous recombination, NADH-dependent fumarate reductase (FRD), phosphoenolpyruvate carboxykinase (PEPCK), covalent flavinylation, Cytb5R domain, reactive oxygen species, positive selection, parasite differentiation

## Abstract

The genome of trypanosomatids is rearranged at the level of repeated sequences, where serve as platforms for amplification or deletion of genomic segments. We report here that the *PEPCK* gene knockout (Δ*pepck*) leads to the selection of such a deletion event between the *FRDg* and *FRDm2* genes to produce a chimeric *FRDg-m2* gene in the Δ*pepck\** cell line. FRDg is expressed in peroxisome-like organelles, named glycosomes, expression of FRDm2 has not been detected to date, and FRDg-m2 is a non-functional cytosolic FRD. Re-expression of FRDg significantly impaired growth of the Δ*pepck\** cells, while inhibition of *FRDg-m2* expression had no effect, which indicated that this recombination event has been selected in the Δ*pepck\** cells to eliminate FRDg. FRD activity was not involved in the FRDg-mediated negative effect, while its auto-flavinylation motif is required to impair growth. Considering that (*i*) FRDs are known to generate reactive oxygen species (ROS) by transferring electrons from their flavin moiety(ies) to oxygen, (*ii*) intracellular ROS production is essential for the differentiation of procyclic to epimastigote forms of the parasite and (*iii*) the fumarate reductase activity is not essential for the parasite, we propose that the main role of FRD is to produce part of the ROS necessary to complete the parasitic cycle in the tsetse fly. In this context, the negative effect of FRDg expression in the PEPCK null background is interpreted as an increased production of ROS from oxygen since fumarate, the natural electron acceptor of FRDg, is no longer produced in glycosomes.

## Introduction

Trypanosomatids, including the human infective *Leishmania* and *Trypanosoma* species, present several biological singularities in comparison with classical eukaryotic model organisms. For instance, genes are transcribed constitutively as part of long polycistronic units where the precursor mRNA molecules are matured by coupled trans-splicing and polyadenylation (1). As a consequence, gene regulation occurs mostly at the post-transcriptional, translational and post-translational levels with no control at the level of transcription initiation. Changes in gene copy number can also modulate gene expression and are therefore seen when selective pressure is applied. They usually arise from homologous recombination events between repeated sequences and are particularly common in *Leishmania* spp. (2). In *Leishmania*, small repetitive sequences are widespread throughout the genome and recombination events appear stochastically with a frequency in the order of 10^−6^/10^−7^ per cell generation. They result either in the production of extrachromosomal DNA sequences or in the deletion of the DNA fragment located between the two recombinogenic repeats. Under selection pressure, such as exposition to drugs, a subpopulation with an advantageous amplicon conferring drug resistance can emerge (3–9). The genome of *T. brucei* also contains a large number of sequence repeats (773) potentially leading to 1848 genetic recombination events, some of them already experimentally validated (2). So far, no DNA amplification (except for changes in ploidy and in gene copy number) has been observed upon specific selection, suggesting that deletions are more common (10–12). We report here the selection of such a stochastic deletion in the genome of *T. brucei* mutants, which is driven by metabolic constraints.

PCF trypanosomes have an elaborate energy metabolism based on glucose or proline, depending on carbon source availability (13). In the glucose-free environment of its insect host (tsetse fly), the parasite depends on proline for its metabolism (14, 15) and needs to produce hexose phosphates through gluconeogenesis from proline-derived phosphoenolpyruvate (PEP) to feed essential pathways (16). Two phosphoenolpyruvate-producing enzymes, PEP carboxykinase (PEPCK, EC: 4.1.1.32, Tb927.2.4210) and pyruvate phosphate dikinase (PPDK, EC 2.7.9.1, Tb927.11.3120) have a redundant function for the essential gluconeogenesis from proline (17). In glucose-rich conditions, PPDK and PEPCK work in the opposite direction to produce pyruvate and PEP, respectively, in addition to ATP. This pathway is also essential to maintain the glycosomal redox balance (18). Glycosomes are specialized peroxisomes, which harbour the 6 or 7 first glycolytic steps (19). Because of the impermeability of the glycosomal membrane to bulky metabolites, such as cofactors and nucleotides, ATP molecules consumed by the first glycolytic steps (steps 1 and 2 in Fig S1) need to be regenerated in the glycosomes by PPDK and PEPCK (step 14 and 15) (18). Similarly, NAD^+^ molecules consumed in the glycosomes during glycolysis (step 6), have to be regenerated within the organelle by the succinic fermentation pathway composed of PEPCK, malate dehydrogenase (EC: 1.1.1.37, Tb927.10.15410, step 16), fumarase (EC: 4.2.1.2, Tb927.10.15410, step 17) and fumarate reductase (FRDg, EC: 1.3.1.6, Tb927.5.930, step 18) (20).

The *T. **brucei*** genome contains three *FRD* genes. Two are tandemly arranged in chromosome 5: they encode the glycosomal isoform (FRDg) and a potential FRD isoform for which expression has not been detected so far in trypanosomes (FRDm2, Tb927.5.940). The third gene, located on chromosome 10, codes for the mitochondrial isoform (FRDm1, Tb927.10.3650, step 21) (21, 22) (Fig 1B). FRDg is a 120 kDa protein composed of three domains, a N-terminal ApbE (<u>A</u>lternative <u>p</u>yrimidine <u>b</u>iosynthesis protein)-like flavin transferase domain (pfam: PF02424, a central FRD domain (superfamily: SSF56425) and a C-terminal cytochrome b5 reductase (Cytb5R) domain (superfamily: SSF63380) (21). In addition FRDg has a conserved flavinylation motif at its extreme N-terminus, shown to be required for FRD activity in the related organism *Leptomonas pyrrhocoris* (23). We report here that two independent PEPCK null mutant cell lines express a chimeric non-functional FRDg-m2 isoform resulting from homologous recombination within the *FRDg*/*FRDm2* locus. The selective advantage provided by the loss of the *FRDg* gene in the context of the PEPCK null background depends on the glycosomal localization of FRDg and the presence of the N-terminal putative flavinylation site. We propose that the absence of metabolic flux through the glycosomal succinic fermentation pathway in PEPCK null mutants made the FAD/FMN cofactors of FRDg available to oxygen for production of reactive oxygen species (ROS) in the organelles.

**Figure 1.**
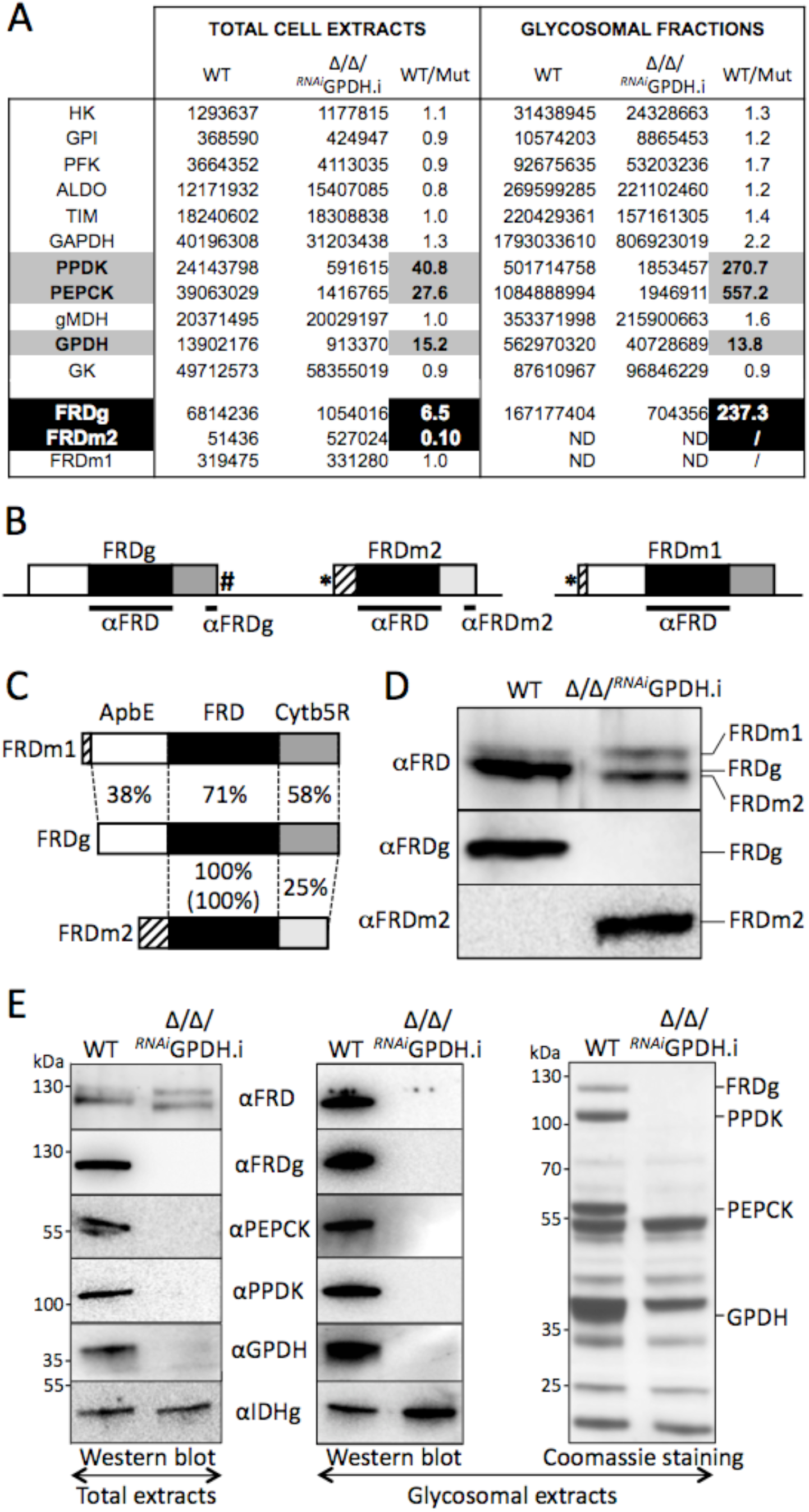
Altered expression of the FRD isoforms in the Δ*ppdk*/Δ*pepck*/^*RNAi*^GPDH mutant cell line. Panel A compares the expression of glycosomal glycolytic enzymes and FRD isoforms obtained by label-free mass spectrometry proteomic analysis (n = 3) of total lysates and glycosomal fractions of the parental (WT) and tetracycline-induced Δ*ppdk*/Δ*pepck*/^*RNAi*^GPDH (Δ/Δ/^*RNAi*^GPDH.i) cell lines (see the PXD020185 dataset in the PRIDE partner repository). The ratio between peptide counts in the parental and mutant cell lines is indicated in the WT/Mut column, with those showing big differences being highlighted. The organization of FRD genes in the *T. brucei* genome is shown in panel B with the glycosomal FRDg and the putative mitochondrial FRDm2 isoforms tandemly arranged on chromosome 5, while the mitochondrial FRDm1 isoform is located on chromosome 10. The white, black and grey (light and dark) boxes represent the ApbE-like, fumarate reductase and cytochrome b5 reductase domains,respectively. Mitochondrial targeting signals present at the N-terminus extremity of FRDm1 (experimentally confirmed (22) and FRDm2 (putative signal corresponding to most of the hatched box) are indicated by asterisks, and the PTS1 motif at the C-terminal end of FRDg is highlighted by a hash. The recombinant protein (aFRD) and peptides (aFRDg and aFRDm2) used for immune sera production are indicated by black bars. Panel C indicates amino acid identity between the 3 domains shared by the FRD isoforms expressed as percentage, with the value into brackets corresponding to nucleotide identity. Expression of the FRD isoforms in the parental (WT) and Δ/Δ/^*RNAi*^GPDH.i cell lines is shown in panel D by Western blot, using antibodies specific for FRDg (aFRDg), FRDm2 (aFRDm2) or the three *T. brucei* FRD isoforms (aFRD). Panel E shows the analysis of the glycosomal localization of the FRD isoforms performed by Western blot with the indicated immune sera on total trypanosome lysates (left panel) and purified glycosomes (central panel) of the parental and Δ*ppdk*/Δ*pepck*/^*RNAi*^GPDH.i cell lines. The glycosomal isocitrate dehydrogenase (IDHg) antibodies were used as a loading control. The right panel is a Coomassie staining of purified glycosomes, which highlights the absence of PPDK, PEPCK, GPDH and FRDg in the glycosomes of the mutant cell line.

## Results

### Expression of a chimeric FRD isoform in the Δppdk/Δpepck/^RNAi^GPDH mutant cell line

In order to study possible changes in gene expression of mutants missing key enzymes involved in the maintenance of the glycosomal redox and ATP/ADP balances, we have compared the total proteomes of the parental, Δ*ppdk* (24), Δ*pepck* (20), Δ*ppdk*/Δ*pepck* (18) and Δ*ppdk*/Δ*pepck*/^*RNAi*^GPDH.i (“.i” stands for tetracycline-induced) cell lines, by label-free quantitative mass spectrometry. The effectiveness of this approach was confirmed by the 19.7- to 41.1-fold reduction observed for the PPDK and/or PEPCK signals in the four mutant cell lines analyzed, compared to the parental cell line (see Fig 1A). Similarly, the GPDH signal was strongly reduced (15.2-fold) in the Δ*ppdk*/Δ*pepck*/^*RNAi*^GPDH.i mutant. This analysis also showed that expression of FRDg and FRDm2 were 6.5-fold decreased and 10-fold increased, respectively, in the Δ*ppdk*/Δ*pepck*/^*RNAi*^GPDH.i cell line, while expression of FRDm1 was not affected (Fig 1A). In contrast, expression of the three FRD isoforms remained unaffected in the three other mutant cell lines (PXD020185 dataset on the ProteomeXchange Consortium). This FRD expression pattern was confirmed by Western blotting using immune sera specific to FRDg (αFRDg) and FRDm2 (αFRDm2), in addition to the αFRD immune serum produced against the conserved FRDg central domain, which is 100% and 71% identical with FRDm2 and FRDm1, respectively (Fig 1B-C). The αFRD antibodies recognized two proteins in both the parental and Δ*ppdk*/Δ*pepck*/^*RNAi*^GPDH.i cell lines, including the ~130 kDa FRDm1 isoform (Fig 1D-E). As previously reported, the second isoform expressed in the parental cell line (~120 kDa) was recognized by the αFRDg, while no signal corresponding to FRDm2 was detected, using αFRDm2 (24). In contrast, the ~115 kDa protein expressed in the Δ*ppdk*/Δ*pepck*/^*RNAi*^GPDH.i cell line was recognized by αFRDm2, but not αFRDg. This suggests that the mutant cell line switched from FRDg to FRDm2 expression, although the apparent size of the detected FRDm2 isoform was higher than the theoretical one (~115 *versus* 94.8 kDa). Coomassie staining, Western blotting (Fig 1E) and proteomic analyses (Fig 1A) of purified glycosomal fractions confirmed the glycosomal localization of FRDg expressed in the parental cell line. In contrast, none of the FRD isoforms were detectable by Western blot in the glycosomal fractions of the mutant cell line, which is consistent with the proteomic analyses.

To determine whether the mutually exclusive expression of FRDg and FRDm2 was related to genomic rearrangement inside the *FRDg*/*FRDm2* locus, a Southern blot analysis was conducted using as probe the conserved FRDg/FRDm2 central domain, which hybridizes with the *FRDm1* gene (“1” in Fig 2A) but gives a much stronger signal for the *FRDg* and *FRDm2* genes (“g” and “2” in Fig 2A). The restriction pattern obtained with the NcoI-, PvuII-, NdeI- and XhoI-digested parental genomic DNA (Fig 2A) was consistent with the restriction map of the FRDm1 (Fig 2B) and FRDg/FRDm2 (Fig 2C) loci deduced from the *T. brucei* TriTrypDB database (strain 927). Although the *FRDm1* locus was identical in the Δ*ppdk*/Δ*pepck*/^*RNAi*^GPDH genome, the pattern observed for the *FRDg*/*FRDm2* locus differed markedly (Fig 2A). For instance, the 6.4 kb PvuII-fragment containing the two FRD genes in the parental genome was converted into a 2.3 kb PvuII-fragment in the mutant genome, suggesting that 4.1 kb had been deleted from the *FRDg/FRDm2* locus. Analysis of the three other restriction profiles led to the same conclusion. The size of the deleted DNA fragment (4.1 kb) was consistent with the size of the theoretical DNA fragment (4074 bp) resulting from homologous recombination between the central 1450 bp FRD domains, which are 100% identical in the *FRDg* and *FRDm2* genes. In conclusion, these data showed that a recombination event occurred between the *FRDg* and *FRDm2* genes in the Δ*ppdk*/Δ*pepck*/^*RNAi*^GPDH mutant to generate a *FRDg-m2* chimeric gene coding for a FRD chimeric protein slightly smaller than FRDg (theoretical molecular weights: 120.6 *versus* 123.5 kDa, respectively). This DNA rearrangement event was present on both alleles of the locus, since the wild-type FRDg and FRDm2 amplified DNA fragments (Fig. 2C) and the endogenous FRDg protein (Fig. 1D) were not detectable in the mutant cell line.

**Figure 2.**
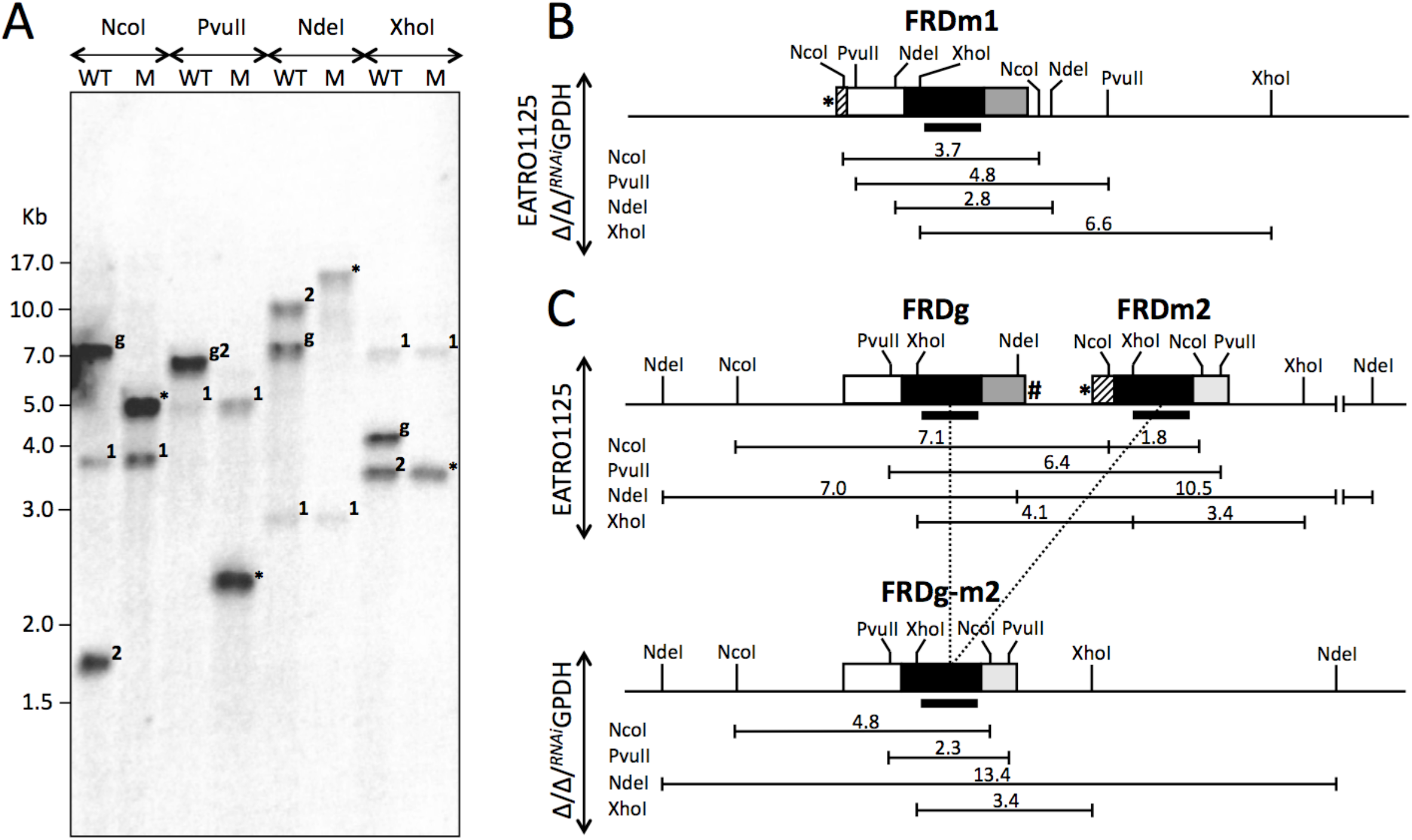
Recombination inside the *FRDg*/*FRDm2* locus in the Δ*ppdk*/Δ*pepck*/^*RNAi*^GPDH cell line. Panel A shows a Southern blot analysis of the parental (WT) and Δ*ppdk*/Δ*pepck*/^*RNAi*^GPDH mutant (M) genomic DNA after digestion with the NcoI, PvuII, NdeI or XhoI restriction enzymes and probing with the *FRDg/FRDm2* central domain, homologous to the *FRDm1* gene (weak signals present in both cell lines). Abbreviations used to identify the labelled fragments: 1, FRDm1; 2, FRDm2; g, FRDg; *, FRDg-m2. The restriction maps presented in panels B (FRDm1) and the upper part of panel C (*FRDg/FRDm2* locus) are deduced from the genome sequence of the 927 strain available in TriTrypDB, while the lower part of panel C represents the FRDg/FRDm2 locus after deletion of the 4.1 kb fragment by homologous recombination (dot lines) in the Δ*ppdk*/Δ*pepck*/^*RNAi*^GPDH mutant cell line (Δ/Δ/^*RNAi*^GPDH). The size of the fragments is indicated in kb and the black bars represent the DNA fragment used to probe the blot. See Fig 1B legend for the color code of the genes.

### The homologous recombination event occurs in wild type cells

To further study this DNA rearrangement event, we used PCR with primer pairs designed for amplification of the central FRD domain of the *FRDg* (g5 and g3 primers), *FRDm2* (m5 and m3 primers) and *FRDg-m2* (g5 and m3 primers) genes (see Fig 3A). As expected, the FRDg- and FRDm2-specific DNA fragments were amplified from the parental EATRO1125 cell line but not from the Δ*ppdk*/Δ*pepck*/^*RNAi*^GPDH genomic DNA (Fig 3B-C), confirming the loss of the wild-type *FRDg/FRDm2* locus in the mutant cell population. Also in agreement with the Southern blot data, the FRDg-m2 specific fragment was amplified from the mutant genomic DNA. Interestingly, however, the FRDg-m2 specific fragment was, also very weakly PCR-amplified from the parental EATRO1125 cell line, which suggests that the recombination event stochastically occurred in the wild-type cells (Fig 3B-C). Moreover, a 1.5kb PCR product was obtained using parental DNA and the g3 and m5 primers; we suggest that the template was a circularized version of the deleted fragment (Fig 3A-C). No corresponding PCR product was detected in the mutant cell line, suggesting that the circularized deleted DNA fragment was not replicated and thus diluted during cell division to become undetectable. The same PCR analysis conducted on genomic DNA samples showed that the rearrangement event occurred in other strains of *Trypanosoma brucei* (*T. equiperdum*, *T. b. brucei* and *T. b. rhodesiense*) (Fig 3B).

**Figure 3.**
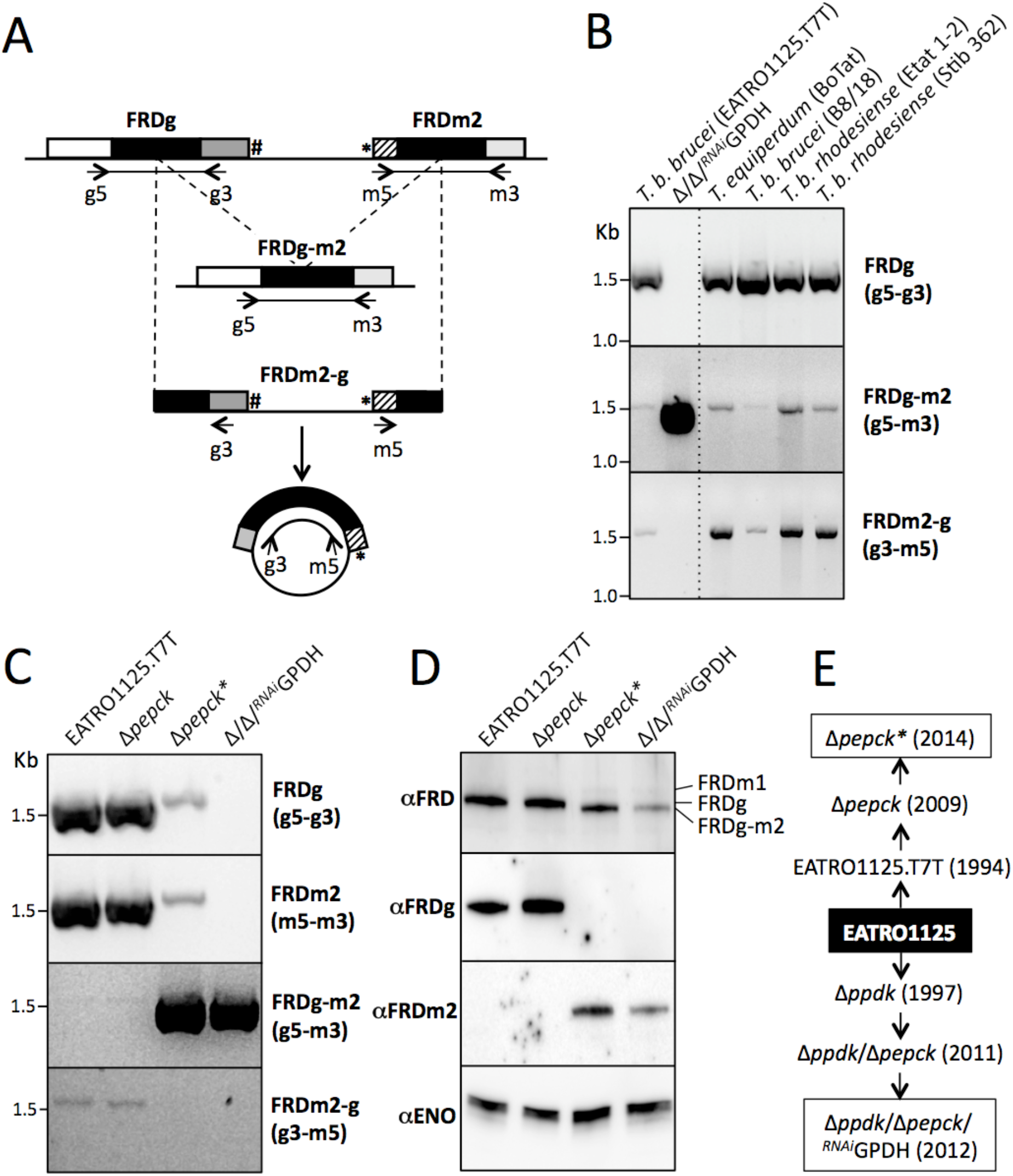
PCR analysis of the recombination event in the *FRDg/FRDm2* locus. The PCR strategy developed to detect a recombination event within the *FRDg/FRDm2* locus is described in panel A. This schematic representation shows the DNA recombination event leading to the deletion of the FRDm2-g fragment, which can be circularized by ligation, as well as the position of the primers (arrows) designed to amplify the FRD domains of *FRDg* (g5-g3; 1583 bp), *FRDm2* (m5-m3; 1611 bp) and the chimeric *FRDg-m2* (g5-m3; 1577 bp), as well as the recircularized deleted FRDm2-g fragment (g3-m5; 1608 bp). See Figure 1B legend for the color code of the genes. Panel B shows a PCR analysis of the EATRO1125.T7T parental and Δ*ppdk*/Δ*pepck*/^*RNAi*^GPDH cell lines, as well as seven additional African trypanosome strains, using 100 ng of genomic DNA (10 μg ml^-1^). In panels C-D, a comparative analysis of the parental EATRO1125.T7T, Δ*pepck*\*, Δ*pepck* and Δ*ppdk*/Δ*pepck*/^*RNAi*^GPDH cell lines is presented, using the PCR approach described in panel A (Panel C) and the Western blot analysis using antibodies specific for Enolase (aENO), FRDg (aFRDg), FRDm2 (aFRDm2) or the three *T. brucei* FRD isoforms (aFRD) (Panel D). The history of the Δ*ppdk*/Δ*pepck*/^*RNAi*^GPDH and Δ*pepck* cell lines (boxed), which have selected the recombinant *FRDg*/*FRDm2* locus, is shown in Panel E. The Δ*pepck* cell line obtained in 2009 contains the parental *FRDg/FRDm2* locus, which became recombined later after long-term in vitro culture (Δ*pepck*\* cell line, 2014).Figure 4

We took advantage of the Δ*ppdk*/Δ*pepck*/^*RNAi*^GPDH cell line being homozygous for the *FRDg-m2* recombinant locus to calculate the allele frequency of *FRDg-m2* in the EATRO1125 parental cell line. We compared the *FRDg-m2* copy number in the two lines by semi-quantitative PCR using different amounts of genomic DNA and the g5-m3 primer pair. Primers specific for a control gene (fructose-1,6-bisphosphatase, Tb927.9.8720) were used for normalisation (Fig 4A). The results showed that the *FRDg-m2* gene copy number is 3,700-times higher in the Δ*ppdk*/Δ*pepck*/^*RNAi*^GPDH homogeneous cell line relative to the heterogeneous parental population (Fig 4B), indicating that at the time of analysis one in 1850 cells in the parental population had a hemizygous recombined allele.

**Figure 4.**
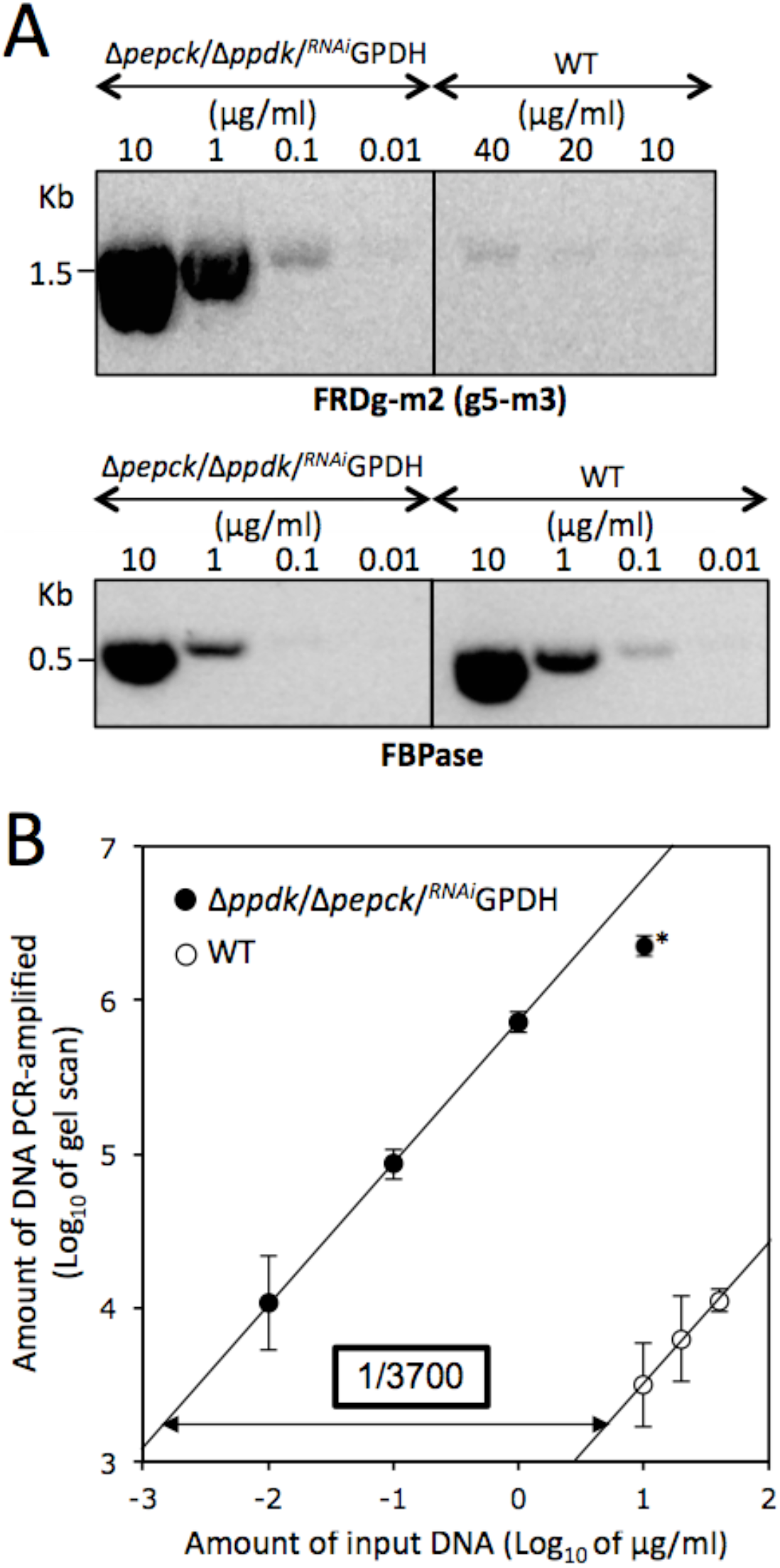
Quantitation of recombined locus frequency. Panel A shows PCR analyses of the wild type (WT) and Δ*ppdk*/Δ*pepck*/^*RNAi*^GPDH cell lines using primers designed for DNA amplification of the chimeric *FRDg-m2* gene (upper panel) or the *FBPase* gene for normalisation (lower panel). The fraction of recombined loci was determined by calculating the relative difference between the two linear regressions of FRDg-m2 PCR signal (gel scan) as a function of the amount of input genomic DNA (panel B). The value obtained for the FRDg-m2 PCR signal with 10 μg ml^-1^ of Δ*ppdk*/Δ*pepck*/^*RNAi*^GPDH genomic DNA, highlighted by an asterisk, was excluded from the trend line.

Altogether, this analysis suggested that the generation of the *FRDg-m2* chimeric gene occurs at low frequency by homologous recombination in the *T. brucei* genome, but was specifically selected in the Δ*ppdk*/Δ*pepck*/^*RNAi*^GPDH cell line.

### Selection of the FRDg-m2 recombinant locus also occurred in the Δpepck mutant cell line

Analysis of the Δ*pepck* mutant obtained and frozen in 2008 (20), then thawed in 2013 and maintained for weeks in culture (see Fig 3E, here named Δ*pepck*\*), yielded more information about selection of the homologous recombination event in the *FRDg*/*FRDm2* locus. It is noteworthy that this Δ*ppdk*/Δ*pepck*/^*RNAi*^GPDH cell line was not derived from the Δ*pepck* cell line (see Fig 3E). Indeed, the Δ*pepck* and the parental cells showed the same PCR profile, while the Δ*pepck\** cells maintained for a long-term in *in vitro* culture showed a pattern similar to the Δ*ppdk*/Δ*pepck*/^*RNAi*^GPDH cell line. Therefore, selection of the recombinant allele occurred independently in a second mutant cell line (Fig 3C). However, the selection process was probably less stringent or more recent in the *PEPCK* null background compared to the Δ*ppdk*/Δ*pepck*/^*RNAi*^GPDH background, as illustrated by the presence of the wild-type locus in the Δ*pepck*\* population (g5-g3 and m5-m3 primer pairs in Fig 3C), even after months of growth. As expected from the PCR analysis, the FRDg-m2 chimeric isoform was expressed in the Δ*pepck*\* cell line, but not detectable by Western blotting in the Δ*pepck* cells (Fig 3D). It is noteworthy that proteomics analysis performed on the Δ*pepck* cell line before long-term cultivation showed an intermediate profile of FRD isoform expression between the parental and the Δ*ppdk*/Δ*pepck*/^*RNAi*^GPDH cell lines (PXD020185 dataset on the ProteomeXchange Consortium). These data strongly suggest selection for the *FRDg-m2* recombinant locus when *PEPCK* is missing (Fig 3E).

### Cytosolic localization of the inactive chimeric FRDg-m2 isoform

The glycosomal localization of FRDg was confirmed by a digitonin cell fractionation experiment. Western blot analysis of the supernatant fractions confirmed that as expected, the FRDg isoform was released together with the PPDK and PEPCK glycosomal markers (Fig 5A). In contrast, the FRDg-m2 chimeric isoform expressed in the Δ*pepck*\* cell line was released at lower digitonin concentrations (0.03 mg *versus* 0.14 mg of digitonin per mg of protein) together with the enolase cytosolic marker (Fig 5A). The cytosolic location of the FRDg-m2 chimeric isoform expressed in the Δ*pepck*\* cell line was confirmed by a Western blot analysis of glycosomal and cytosolic fractions prepared by differential centrifugation after silicon carbide cell homogenization (Fig 5B).

**Figure 5.**
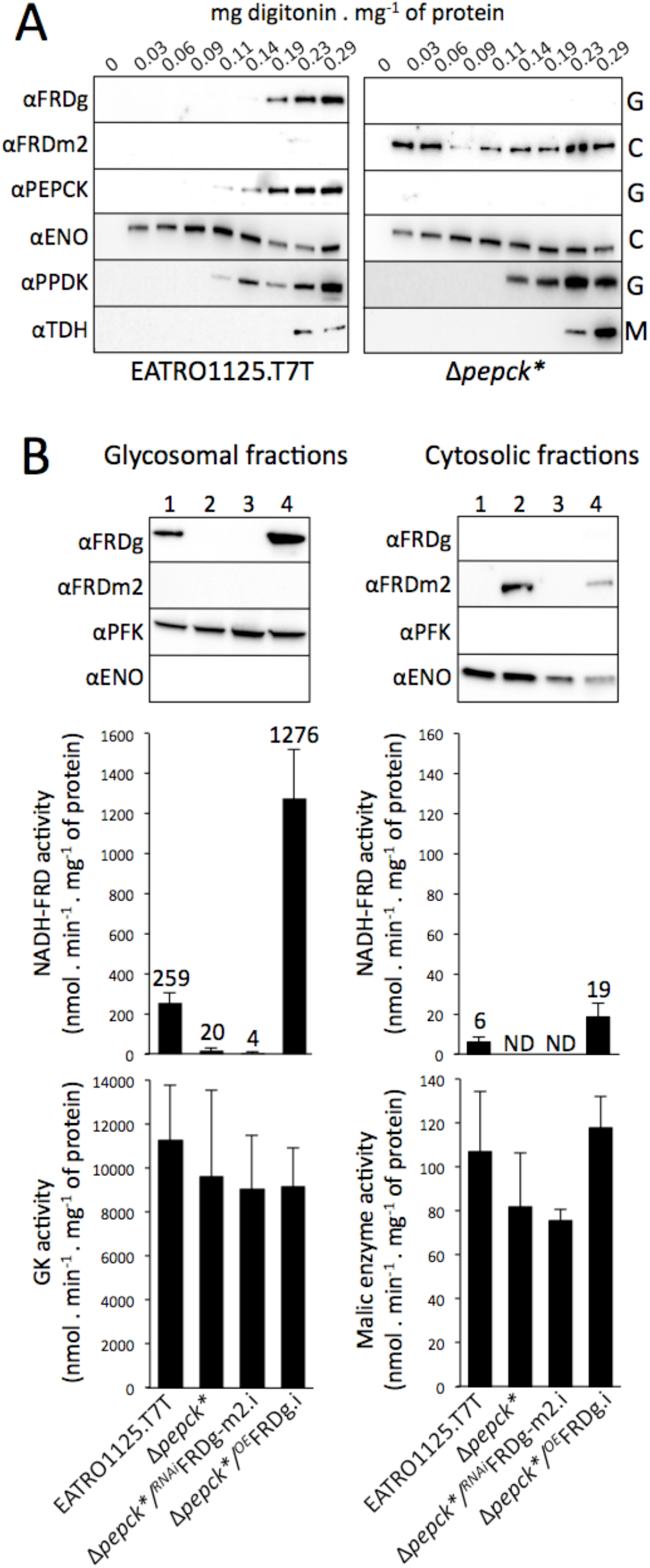
The cytosolic FRDg-m2 chimeric isoform is not enzymatically active. Panel A shows the glycosomal and cytosolic localization of FRDg and the chimeric FRDg-m2 isoforms, respectively, by digitonin titration. The supernatant collected from the EATRO1125.T7T and Δ*pepck*\* cells incubated with 0-0.29 mg of digitonin per mg of protein was analyzed by Western blot using the anti-FRDg, anti-FRDm2 as well as immune sera against cytosolic (enolase, ENO), glycosomal (PPDK) and mitochondrial (threonine dehydrogenase, TDH) markers. Panel B shows the Western blotting and enzymatic activities determined in the glycosomal and cytosolic fractions of EATRO1125.T7T (1), Δ*pepck*\* (2), Δ*pepck*\*/^*RNAi*^FRDg-m2.i (3) and Δ*pepck*\*/^*OE*^FRDg.i (4) cells lines. Expression of FRDg and the chimeric FRDg-m2 isoforms was determined by Western blotting using the anti-FRDg and anti-FRDm2 immune sera (top panel). Immune sera against the glycosomal phosphofructokinase (PFK) and the cytosol enolase (ENO) were used as loading controls. NADH-FRD activity was determined on the same fractions used for Western blot analyses. For normalization, the glycerol kinase (GK) and malic enzyme activities were also determined in the glycosomal and cytosolic fractions (lower panel).

The NADH-dependent FRD (NADH-FRD) activity was determined in the glycosomal and cytosolic fractions of the original EATRO1125.T7T and the Δ*pepck*\* cell lines. As expected, NADH-FRD activity was detected in the glycosomal fraction of cells expressing FRDg (EATRO1125.T7T), but not in the glycosomes of the Δ*pepck\** cell line (Fig 5B). The low level of NADH-FRD activity detected in the cytosolic fraction of the EATRO1125.T7T cell line, compared to the glycosomal fraction (2.3%), was presumably due to the lysis of a few glycosomes during the grinding step. The absence of NADH-FRD activity in the cytosolic fraction of the Δ*pepck*\* cell line demonstrates that the chimeric FRDg-m2 isoform is inactive (Fig 5B). These data highlight the role of the Cytb5R domain in the NADH-FRD activity, since only this domain differs between the active FRDg and inactive FRDg-m2 isoforms.

### Expression of FRDg affects the Δpepck* growth rate

Selection of the *FRDg-m2* recombinant locus in the Δ*pepck\** and Δ*ppdk*/Δ*pepck*/^*RNAi*^GPDH cell lines implied that either expression of the FRDg-m2 isoform in the cytosol, or abolition of FRDg expression in the glycosomes, provided a selective advantage to both mutant cell lines. To determine which of these two hypotheses is correct, tetracycline-inducible ectopic expression of FRDg and RNAi-mediated down-regulation of FRDg-m2 were performed in the Δ*pepck\** cell line (Δ*pepck\**/^*OE*^FRDg and Δ*pepck\**/^*RNAi*^FRDg-m2, respectively) (Fig 6A). These experiments could not be conducted with the Δ*ppdk*/Δ*pepck*/^*RNAi*^GPDH cell line, because all five available selectable markers had already been used. The glycosomal localization of the recombinant FRDg in the Δ*pepck\**/^*OE*^FRDg.i line was confirmed by Western blotting and enzymatic activity assay of glycosomal fractions (Fig 5B). The doubling time of the Δ*pepck*\*/^*RNAi*^FRDg-m2 cell population was identical in the absence (.ni) or the presence (.i) of tetracycline, indicating that expression of the FRDg-m2 chimera was well tolerated by the Δ*pepck*\* mutant. In contrast, induction of FRDg expression induced a slightly reduced the growth rate of the Δ*pepck*\*/^*OE*^FRDg.i cell line (Fig 6B).

**Figure 6.**
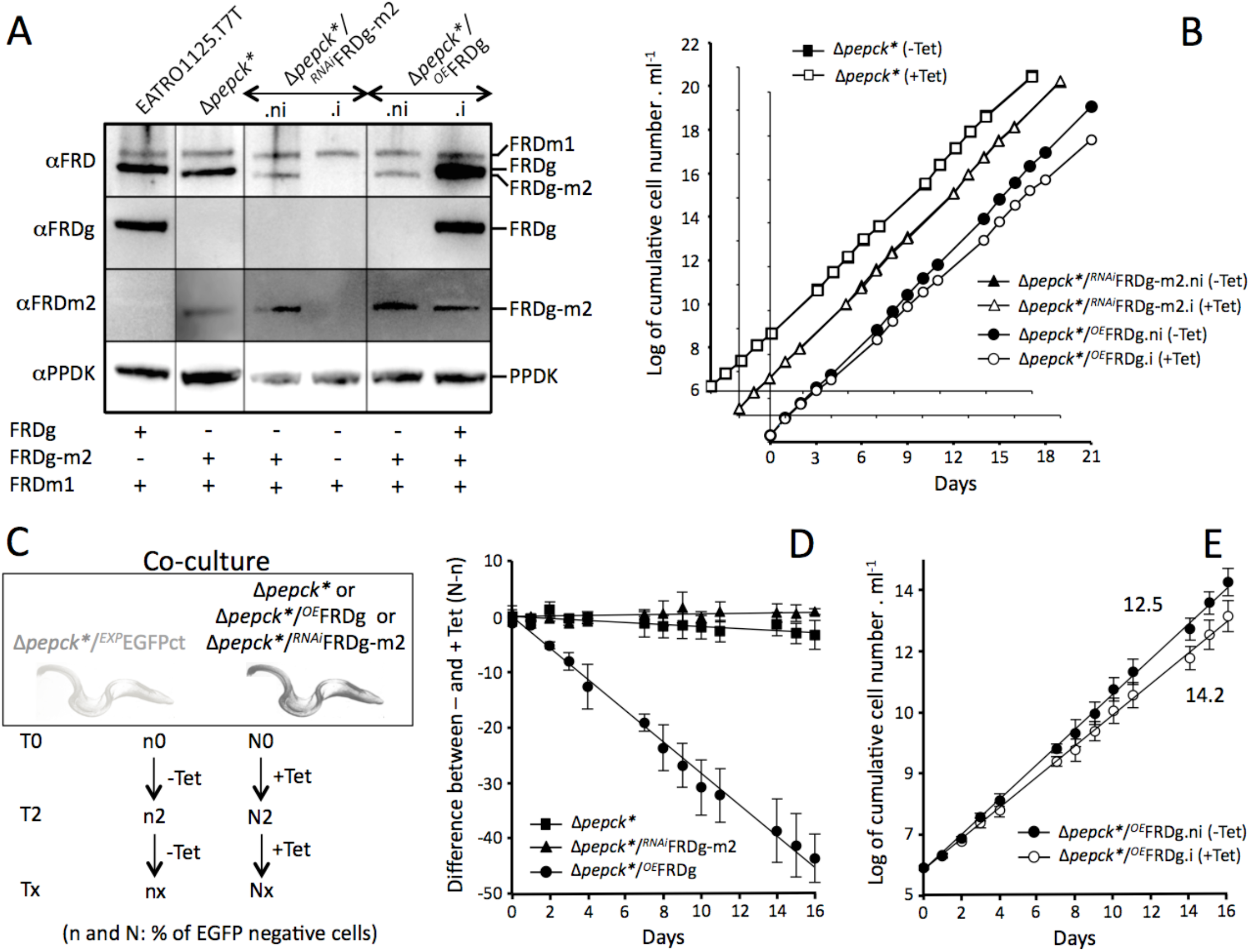
Expression of FRDg is responsible for reduced growth of the Δ*pepck*\* cell line. In panel A, expression of the FRD isoforms in the EATRO1125.T7T and Δ*pepck*\* parental cell lines, as well as the tetracycline-induced (.i) or non-induced (.ni) Δ*pepck*\*/^*RNAi*^FRDg-m2 and Δ*pepck*\*/^*OE*^FRDg was monitored by Western blot analysis using immune sera indicated on the left margin. Expression of the FRD isoform(s) is indicated under the blot. The growth curves of these tetracycline-induced (.i or +Tet) or non-induced (.ni or −Tet) cell lines are shown in panel B. To confirm the moderate growth defect observed for the Δ*pepck*\* mutant expressing the recombinant FRDg (Δ*pepck*\*/^*OE*^FRDg.i), the Δ*pepck*\*, Δ*pepck*\*/^*RNAi*^FRDg-m2 and Δ*pepck*\*/^*OE*^FRDg cell lines were co-cultured with the Δ*pepck*\* cell line constitutively expressing EGFP (Δ*pepck*\*/^*OE*^EGFPct), in the presence or the absence of tetracycline. Flow cytometry analyses were conducted to determine EGFP positive cells (Δ*pepck*\*/^*OE*^EGFPct) and EGFP negative cells (Δ*pepck*\* or double mutant cell lines) all along the growth curve, as illustrated in panel C. Panel D shows the difference of the percentage of EGFP negative cells between non-induced and induced conditions, all along the 16-day co-culture (mean of 3 independent experiments). The growth curve of induced and non-induced Δ*pepck*\*/^*OE*^FRDg co-cultured with the EGFP-tagged parental cell line (Δ*pepck*\*/^*OE*^EGFPct) was deduced from the same datasets and plotted in panel E. The numbers indicate the population doubling time in both growth conditions.

Selection against FRDg expression was confirmed in a co-culture experiment. EGFP-tagged Δ*pepck*\* cell line (Δ*pepck*\*/^*OE*^EGFPct, constitutively expressing EGFP) was co-cultured with the Δ*pepck*\*(control), Δ*pepck*\*/^*RNAi*^FRDg-m2 or Δ*pepck*\*/^*OE*^FRDg cell lines in the presence or the absence of tetracycline, and the proportion of EGFP-positive cells was determined over time by flow cytometry (Fig 6C). The percentage of EGFP negative cells between induced and non-induced conditions gradually decreased when FRDg was expressed in the Δ*pepck*\* cell line, while there was no selection against FRDg-m2 expression (Fig 6D). Calculations revealed that expression of FRDg increased the doubling time of the Δ*pepck*\* cells from 12.5 h to 14.2 h (Fig 6E).

Recombinant FRDg was ~4.5-times more expressed in the Δ*pepck*\*/^*OE*^FRDg.i cell line than the endogenous FRDg in the parental WT cells (Fig 5B). Interestingly, overexpression of FRDg in the WT background (^*OE*^FRDg.i) also induced a significant growth defect (Fig 7A). FRDg overexpression is therefore detrimental in the absence as well as in the presence of PEPCK. This does not, however, affect our overall conclusions: the native levels of FRDg expression must affect the Δ*pepck* and Δ*ppdk*/Δ*pepck*/^*RNAi*^GPDH cell lines more than the EATRO1125.T7T parental cell line, since the recombined *FRDg*/*FRDm2* locus has been positively selected in the mutant cell lines.

**Figure 7.**
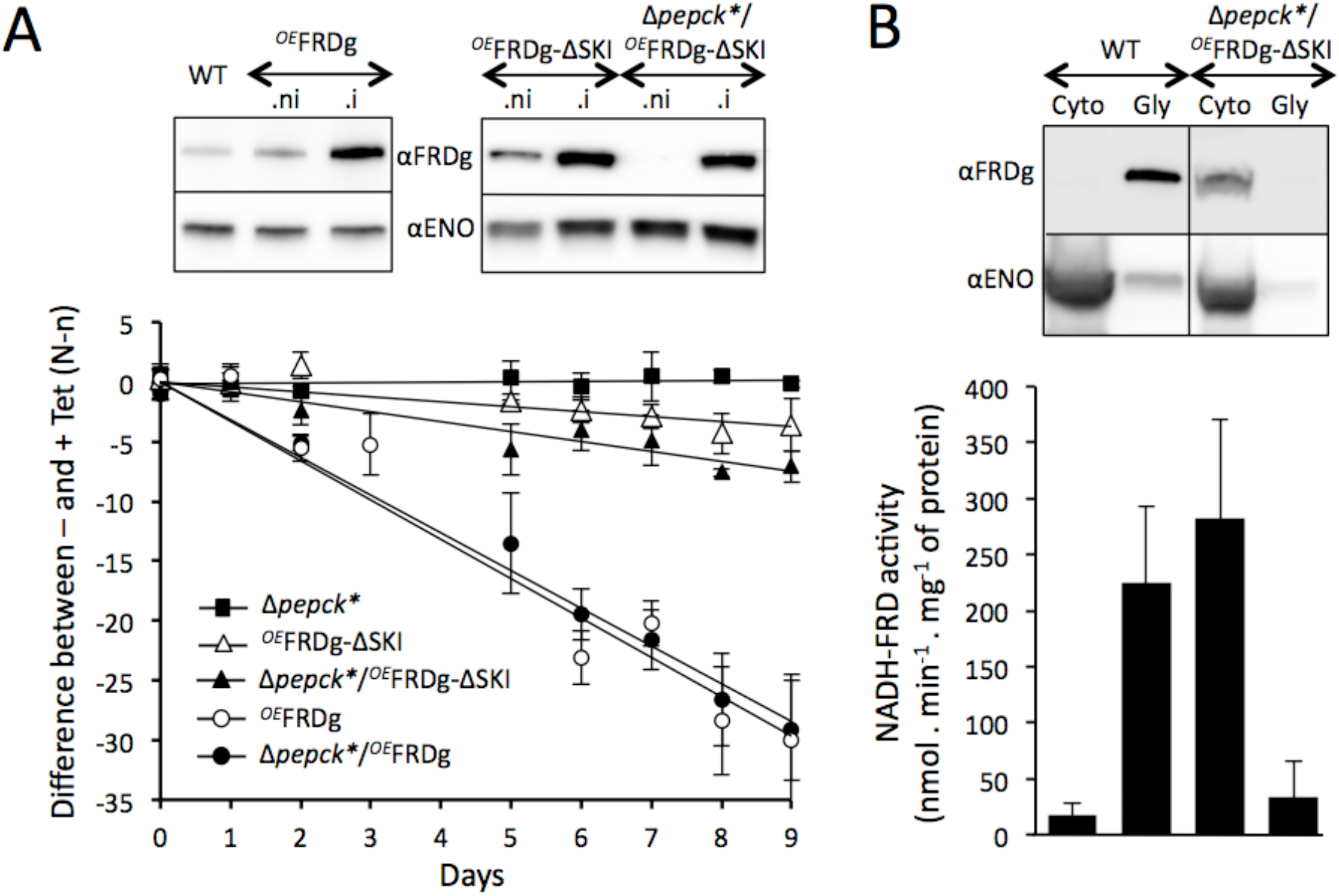
The cytosolic expression of FRDg does not affect growth of the Δ*pepck*\* cell line. Panel A shows the effect of the expression of full length FRDg and FRDg-ΔSKI in the WT (^*OE*^FRDg and *OE FRDg-*ΔSKI) or the Δ*pepck*\* background (Δ*pepck*\*/^*OE*^FRDg and Δ*pepck*\*/^*OE*^*FRDg-*ΔSKI) using as negative and positive controls the Δ*pepck*\* cell line. For this experiment, the mutant cell lines were co-cultured with the Δ*pepck*\* cell line constitutively expressing EGFP (Δ*pepck*\*/^*OE*^EGFPct), in the presence or the absence of tetracycline, as described in Figure 6C. The difference of the percentage of EGFP negative cells between non-induced and induced conditions is plotted as a function of time of growth (mean of 3 independent experiments). The top panel shows a Western blot analysis of these tetracycline-induced (.i) or non-induced (.ni) cell lines. In panel B, expression of the endogenous FRDg isoform in the parental EATRO1125.T7T cell line (WT) or the recombinant *FRDg-*ΔSKI in the tetracycline-induced Δ*pepck*\*/^*OE*^FRDg-ΔSKI mutant was monitored in the glycosomal (Gly) and cytosolic (Cyto) fractions by Western blot analysis using the immune sera indicated on the left margin.The lower panel shows the glycosomal and cytosolic NADH-FRD activities normalized with the GK and malic enzyme activities respectively.

### The glycosomal localization of FRDg is required for the growth retardation

We next tested whether the negative effect of FRDg expression in the Δ*pepck\** background was dependent on the glycosomal localization of the protein. To address this question, the C-terminal peroxisomal targeting signal (PTS1) composed of the last 3 amino acids, the SKI tripeptide in FRDg, was removed from the recombinant protein to express a functional FRD in the cytosol of the Δ*pepck*\* cell line (Δ*pepck*\*/^*OE*^FRDg-ΔSKI). The cytosolic localization of the FRDg-ΔSKI was confirmed by Western blotting of the glycosomal and cytosolic fractions (Fig 7A), and NADH-FRD activity was detected in the cytosolic fraction of the Δ*pepck*\*/^*OE*^FRDg-ΔSKI.i cell line (Fig 5B). Co-culture of the Δ*pepck*\*/^*OE*^EGFPct cell line with the Δ*pepck*\*/^*OE*^FRDg-ΔSKI or ^*OE*^FRDg-ΔSKI mutants in the presence or the absence of tetracycline, performed as above, revealed only minimal growth retardation upon FRDg-ΔSKI expression in the Δ*pepck*\* or WT backgrounds (Fig 7B). Thus the glycosomal expression of FRDg was required to affect growth of the Δ*pepck*\* cell line.

### The glycosomal NADH-FRD activity is not responsible for the growth retardation

FRDg is composed of a N-terminal ApbE-like, a central fumarate reductase (FRD) and a C-terminal cytochrome b5 reductase domain (Cytb5R) (Fig 1B). To determine the FRDg domains responsible for the negative growth effect of the FRDg in the Δ*pepck* background, truncated recombinant FRDg proteins missing the FRD (Δctl), ApbE-like (ΔNterm) or ApbE-like/FRD (ΔNterm/ctl) domains were expressed in the Δ*pepck*\* cell line (Δ*pepck\**/^*OE*^FRDg-Δctl, Δ*pepck\**/^*OE*^FRDg-ΔNterm and Δ*pepck\**/^*OE*^FRDg-ΔNterm/ctl, respectively). All three recombinant FRDg proteins were successfully expressed in the glycosomes, and as expected none of them showed NADH-FRD activity (Fig 8A). The Cytb5R domain alone (FRDg-ΔNterm/ctl) or in combination with the FRD domain (FRDg-ΔNterm) did not affect growth, but surprisingly, expression of FRDg-Δctl was even more deleterious than full-length, FRDg (Fig 8B). Clearly, specific NADH-FRD enzymatic activity was not responsible for growth retardation (Fig 8B). Overexpression of recombinant ^*OE*^FRDg-Δctl also inhibited growth of the EATRO1125.T7T parental cell line.

**Figure 8.**
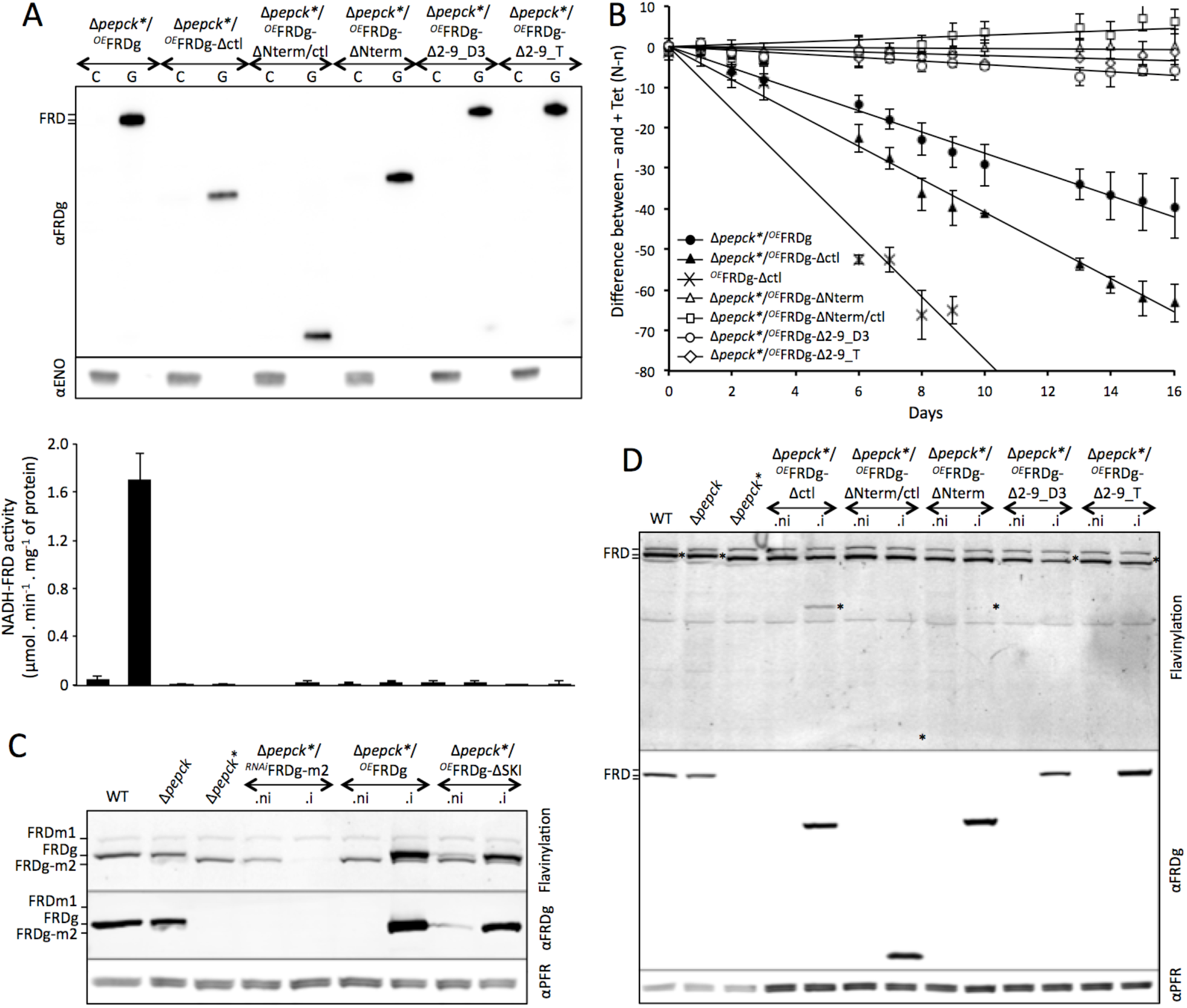
The flavinylation motif of the FRDg N-terminal domain is required for growth retardation of the Δ*pepck*\* cell line. Panel A shows Western blot analyses of cytosol-enriched (C) and glycosome-enriched (G) fractions from Δ*pepck\** cell lines expressing truncated or mutated recombinant FRDg. See Figure 5 for the immune sera used. The lower panel shows the glycosomal and cytosolic NADH-FRD activities normalized with the GK and malic enzyme activities respectively. Panel B shows the effect of the expression of FRDg, FRDg-Δctl, FRDg-ΔNterm, FRDg-ΔNterm/ctl and FRDg-Δ2-9 (clones D3 and T) in the Δ*pepck*\* background and expression of FRDg-Δctl in the parental EATRO1125.T7T (WT) background, as described in Figures 6 and 7. The Western blot control of the ^*OE*^FRDg-Δctl cell line is shown in Figure S2. The difference of the percentage of EGFP negative cells between non-induced and induced conditions is plotted as a function of time of growth (mean of 3 independent experiments). Panels C and D show the presence of covalent flavinylation of the FRD isoforms and mutants in the cell lines analyzed (.i, tetracycline-induced; .ni, non-induced. The top panels show directly detected fluorescence of covalently bound flavin on a denaturing gel, while the lower panels show Western blot analyses with the anti-FRDg and anti-PFR (internal loading reference) immune sera. The locations of the endogenous and recombinant FRDg protein bands are indicated by an asterisk (*) on the direct fluorescence gel image; the positions of FRD isoform bands are indicated at the left gel margin, as detailed in C.

### Growth retardation depends on flavinylation

The absence of an altered growth phenotype upon expression of FRDg-ΔNterm suggested a possible role of the ApbE-like domain. Recently, Serebryakova *et al.* showed that the orthologous FRDg of *Leptomonas pyrrhocoris*, a trypanosomatid related to trypanosomes, contains a covalently attached flavin at serine 9 of the N-terminal flavinylation motif [D_3_(g/s)=(s/t)(s/g)AS_9_]. They suggested that the ApbE-like domain may catalyze the transfer of FMN from FAD to serine 9 of FRDg. Replacement of S9 by an asparagine residue abolished both flavinylation and NADH-fumarate reductase activity of *Leptomonas* FRDg (23). We therefore addressed the role of flavinylation for FRD activity in trypanosomes and for the growth phenotype caused by glycosomal expression of FRDg in the Δ*pepck\** background. The FRDg-Δ2-9 mutant protein missing the first nine N-terminal residues (the suggested flavinylation motif), expressed in the Δ*pepck\** background (Δ*pepck\**/^*OE*^FRDg-Δ2-9_D3 and Δ*pepck\**/^*OE*^FRDg-Δ2-9_T cell lines) did not confer glycosomal FRD activity (Fig 8A) upon tetracycline-induced expression and caused no growth retardation (Fig 8B).

The flavinylation of all endogenous and expressed FRD isoforms and mutants was directly assessed by in gel detection of flavin fluorescence at 526 nm. Using denaturing gels and boiling of protein samples, only covalently linked flavin was detected (Fig 8C-D). For the glycosomally expressed FRD mutant proteins, flavinylation correlates with growth phenotype (Fig 9). This confirmed that covalent flavinylation is required for FRD activity *in vivo* in trypanosomes (Fig 8A). The essential role of FRD flavinylation for the altered growth phenotype implicates electron transfer, albeit not to fumarate, in the deleterious effects.

**Figure 9.**
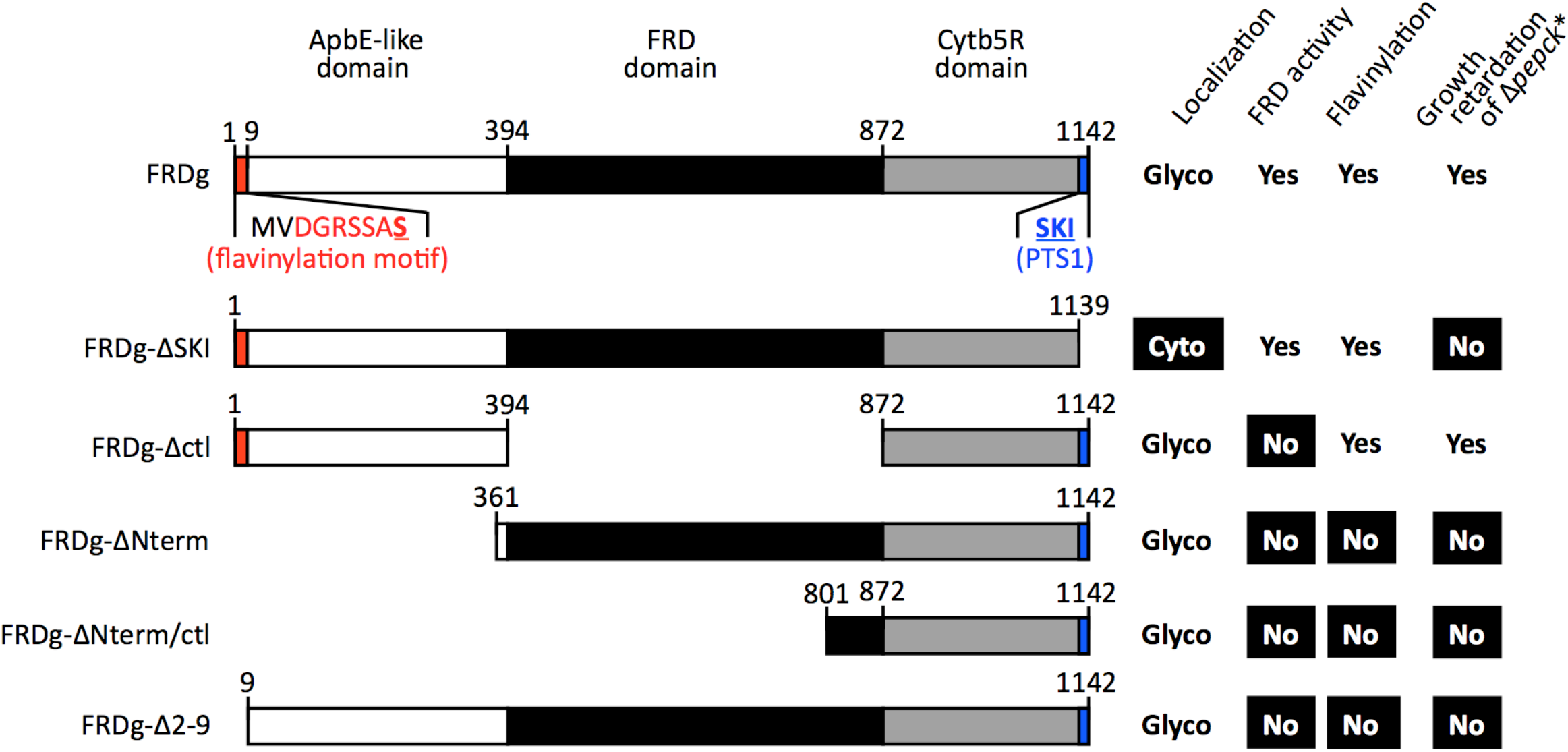
Correlations between FRD domains, FRD flavinylation, glycosomal localization and effect on growth of the Δ*pepck*\* cell line. This figure summarizes expression of the endogenous and mutated FRDg in the Δ*pepck*\* background, their subcellular localization, NADH-dependent FRD activity, covalent flavinylation and the effect on growth of the Δ*pepck*\* cell line. The white, black and grey boxes represent the ApbE-like, fumarate reductase and cytochrome *b*_5_ reductase domains, respectively, and the consensus flavinylation motif is indicated in red (the serine residue that is most likely the covalent attachment site of the flavin moiety is bold and underlined). Difference of phenotype compared to the Δ*pepck*\*/^*OE*^FRDg (FRDg) cell line expressing endogenous FRDg is indicated in white on a black background. Glyco, glycosome; Cyto, cytosol.

### Absence of significant FRDg-catalyzed flux in the Δpepck mutant

Why does FRDg impair growth of the Δ*pepck* cell line? FAD containing enzymes, including FRD, are known to transfer electrons very efficiently to oxygen to generate toxic reactive oxygen species (ROS). We therefore hypothesized that in the absence of metabolic flux through the glycosomal succinate branch, the decrease of fumarate allows oxygen to become the main FRDg substrate. To determine the contribution of FRDg to the production of succinate, we analyzed excreted end products (exometabolome) from the metabolism of the two main carbon sources used by the parental and mutant cell lines (glucose and proline) using the 1H-NMR profiling approach. The advantage of ^1^H-NMR spectrometry is the possibility to distinguish protons bound to ^12^C and ^13^C carbons, so that end products excreted from two different carbon sources can be distinguished, provided that one is uniformly ^13^C-enriched. To achieve this, we used U-^13^C- proline (25). The parental trypanosomes incubated in the presence of glucose and proline produced 71% acetate, 18% succinate and 11% alanine, as excreted end products (Table 1). As expected, succinate was no longer produced from glucose in the Δ*pepck* cell line (20), while the amounts of proline-derived succinate were not affected (see Fig S1). The metabolic patterns of the Δ*pepck* and Δ*pepck\** cell lines were similar, indicating that succinate excreted from proline in the PEPCK null background was produced not in the glycosomes, but in the mitochondrion by succinyl-CoA synthetase as previously proposed (25). Our interpretation of these data is consistent with the maintenance of the level of succinate production from proline in the Δ*pepck\**/^*OE*^FRDg.i mutant, in which expression of FRDg is 4.5 times higher than in the EATRO1125.T7T cell line (WT) (Table 1). As expected, no difference was also observed in the Δ*pepck\**/^*RNAi*^FRDg-m2.i (Table 1). These data suggest that the observed growth phenotype is probably the consequence of the absence of succinate production within the glycosomes, which may lead to an increase production of ROS by FRDg.

**Table 1.** Excreted end products of glucose and glycerol metabolism by BSF parental and mutant cell lines.

|  | WT | *Δpepck* | *Δpepck\** | *Δpepck\*/*<br>*OE FRDg.ni*<sup>a</sup> | *Δpepck\*/*<br>*OE FRDg.i*<sup>a</sup> | *Δpepck\*/*<br>*RNAi FRDg-*<br>*m2.ni* | *Δpepck\*/*<br>*RNAi FRDg-*<br>*m2.i* |
| --- | --- | --- | --- | --- | --- | --- | --- |
|  | nmol.h<sup>-1</sup>.10<sup>8</sup> cells<sup>-1</sup><sup>b</sup> |  |  |  |  |  |  |
| n<sup>c</sup> | 3 | 3 | 3 | 3 | 3 | 3 | 3 |
| Succinate (glucose<sup>d</sup>) | 277 ±15 | nd<sup>e</sup> | nd | nd | nd | nd | nd |
| Succinate (proline<sup>d</sup>) | 132 ±19 | 147 ±21 | 138 ±14 | 94 ±7 | 78 ±6 | 78 ±4 | 72 ±4 |
| Acetate (glucose) | 1431 ±121 | 385 ±25 | 400 ±39 | 414 ±32 | 409 ±10 | 402 ±20 | 394 ±11 |
| Acetate (proline) | 238 ±12 | 366 ±26 | 395 ±39 | 308 ±16 | 330 ±18 | 345 ±19 | 347 ±15 |
| Alanine (glucose) | 229 ±25 | 374 ±29 | 405 ±41 | 367 ±23 | 391 ±13 | 409 ±14 | 400 ±7 |
| Alanine (proline) | 31 ±4 | 142 ±11 | 156 ±20 | 163 ±8 | 182 ±15 | 140 ±15 | 141 ±8 |
| **Total (glucose)** | **1950 ±120** | **759 ±54** | **806 ±78** | **781 ±54** | **800 ±12** | **811 ±34** | **794 ±16** |
| **Total (proline)** | **401 ±15** | **655 ±56** | **689 ±71** | **565 ±29** | **590 ±15** | **563 ±28** | **561 ±20** |

<sup>a</sup> .i: RNAi cell line induced during 5 days by addition of tetracycline; .ni: non-induced RNAi cell line
<sup>b</sup> The amounts of end products excreted from glucose and proline metabolism are expressed as nmoles excreted per hour and per 10<sup>8</sup> cells
<sup>c</sup> Number of biological replicates
<sup>d</sup> Carbon source metabolized into succinate
<sup>e</sup> nd: not detectable

## Discussion

Stochastic recombination leading to amplification of chromosomal regions located between homologous direct or inverted repeated sequences has been observed in *Leishmania major* (2). This genome-wide phenomenon leads to extrachromosomal circular or linear amplified DNAs, as well as deletion of DNA fragments. DNA amplification through stochastic recombination has a direct impact on gene dosage and fosters the selection of adaptive traits in response to environmental pressure, such as drug exposure, as previously reported on many occasions (26). However, the benefit provided by deletion of DNA fragments is much less obvious. In contrast to *Leishmania* spp., the involvement of stochastic recombination in adaptation to environmental pressure has not been reported so far for *T. brucei*, presumably due to a stricter replication system where episomal vector maintenance is the exception (27). This implies that gene deletions or generation of mosaic genes are the observable effects of stochastic recombination in *T. brucei* (11, 28, 29). Δ*pepck* cell lines expressing the chimeric FRDg-m2 instead of FRDg constitute one of the two examples of a stochastic recombination event providing a selective advantage to *T. brucei* reported to date. The other example has been described in the tandemly arranged genes encoding aquaglyceroporins (AQP2 and AQP3), which facilitate the transmembrane transport of water and small nonionic solutes. In *T. brucei*, AQP3 plays a role in the osmoregulation and transport of glycerol, while AQP2 is also a carrier involved in importing the trypanocidal melarsoprol and pentamidine drugs by endocytosis (11, 30, 31). In some melarsoprol/pentamidine resistant cell lines the *AQP2* and *AQP3* genes are replaced by a chimeric *AQP2/3* gene, which lost the capacity to interact with the drugs (11, 29). Since the *AQP2* (Tb927.10.14170) and *AQP3* (Tb927.10.14180) genes are 80% identical, with three ~80 bp direct repeats, and are only separated by 857 bp of non coding sequence, formation of the *AQP2/3* chimeric gene most probably results from drug selection of stochastic homologous recombination within the *AQP2*/*AQP3* locus, as described here for the *FRDg*/*FRDm2* locus.

The gene encoding FRDg is composed of three domains, the central FRD domain (433 aa) flanked by the C-terminal Cytb5R domain (222 aa) and the N-terminal ApbE-like flavin transferase domain (287 aa) (21–23). FRDg is responsible for the glycosomal NADH-dependent FRD activity, however the role of the N- and C-terminal domains is currently unknown. Here, we provide evidence that the Cytb5R domain is required for the FRD activity, since the FRDg-m2 chimera expressed in the cytosol of the Δ*pepck* cell line is not active. Indeed, the only difference between the endogenous FRDg and chimeric FRDg-m2 isoforms is the C-terminal domain showing 34% (e-value: 9 × 10^−32^) and 27% (evalue: 3 × 10^−8^) identity with the *Cryptococcus neoformans* Cytb5R, respectively. Since, the NADH-cytochrome b5 reductases are known to transfer electrons from NADH to cytochrome b5 (32), one may consider that the Cytb5R domain is part of the electron transfer channel required for the FRD activity. Elucidating the 3D structure of FRDg is required to define the mechanism of FRDg activity and characterize this electron transfer channel.

Our data suggest that the FRDm2 isoform has lost FRD activity, since it shares the C-terminal domain of the inactive FRDg-m2 chimera and more importantly it lacks the N-terminal ApbE-like domain including the flavinylation motif (see Fig 1C, (22). The *FRDg*/*FRDm2* locus is conserved across the trypanosomatid lineage, with the FRD/Cytb5R composite structure of the FRDm2 isoforms conserved within the *Trypanosoma* species, while *Crithidia fasciculata* and the *Leishmania* spp. have lost the C-terminal Cytb5R domain. The conservation of FRDm2 in the *Trypanosoma* and *Leishmania* branches, which separated 400-600 millions years ago (33) is surprising as expression of the FRDm2 isoform was neither detectable in the PCF nor in bloodstream forms of *T. brucei* (22) and absence of FRD-specific activity, leaving a biological function enigmatic. Perhaps FRDm2 has a role in stages that have yet to be examined in detail, such as the epimastigotes.

Selection of the FRDg-m2 chimera in the Δ*pepck* and Δ*ppdk*/Δ*pepck*/^*RNAi*^GPDH cell lines implies that this stochastic recombination event was beneficial for the PCF trypanosomes in the context of the PEPCK null background. We demonstrated that this selection is driven by the deleterious effect of FRDg overexpression in the PEPCK null background, which provided a rational explanation for the selection of the recombined Δ*pepck\** cells. It is noteworthy that a ~5-fold overexpression of FRDg also slightly affected growth of the parental EATRO1125.T7T cells, suggesting that the normal FRDg level is compatible with - and perhaps supports - optimal growth of the wild-type parasite, whereas it impairs growth of the Δ*pepck* mutant. This hypothesis may explain why the recombination event has been selected in the PEPCK null background but not in the wild-type background.

How to explain the growth phenotype due to overexpression of FRDg? Several observations support the view that FRDg, as already observed for FRD from other organisms, can produce reactive oxygen species (ROS), known to be toxic at high concentrations. For instance, FRD is a major contributor to ROS formation in *Bacteroides fragilis* exposed to oxygen (34). ROS are formed by autoxidation when redox enzymes accidentally transfer electrons to oxygen rather than to their physiological substrates. This enzymatic promiscuity is well illustrated by the *E. coli* aspartate:fumarate oxidoreductase which was conventionally named aspartate oxidase since oxygen is used as electron acceptor in aerobic conditions (35). However, this enzyme can also transfer electrons to fumarate, which is certainly its, natural substrate in the anaerobic conditions encountered in the intestine by *E. coli* (36). This autoxidation activity of FAD-dependent redox enzymes is due to the solvent accessibility of the flavin moiety, which is situated at the protein surface in order to interact with soluble substrates, as described for the *E. coli* FRD in the absence of its natural substrate, *i.e.* fumarate (37). The notion that oxygen and fumarate compete for electrons provided by FAD was also reported for the *T. brucei* FRD since fumarate inhibited hydrogen peroxide formation with the same affinity as it stimulated NADH-dependent FRD activity (K_*i*_ = 16 *versus* 20 μM) (38). Thus, abolition of glycosomal succinate production in the Δ*pepck* background, which is probably due to a significant reduction of the glycosomal amounts of fumarate (see Fig S1), might stimulate autoxidation activity of FRDg (Fig 10). This hypothesis is consistent with the absence of growth phenotype for the Δ*pepck\**/^*OE*^FRDg-Δ2-9 cell lines, which lost the covalently bound flavin required to transfer electron to the acceptor (Fig 8B). More importantly, the significant increase of the doubling time caused by overexpression of the recombinant FRDg-Δctl in Δ*pepck\** backgrounds, as well as in the wild-type cells, is consistent with the proposed competition between fumarate and oxygen for electrons provided by the covalently bound flavin in the ApbE domain. Indeed, in the absence of the central FRD domain, oxygen would become the main electron acceptor, regardless of the amounts of fumarate inside the glycosomes.

**Figure 10.**
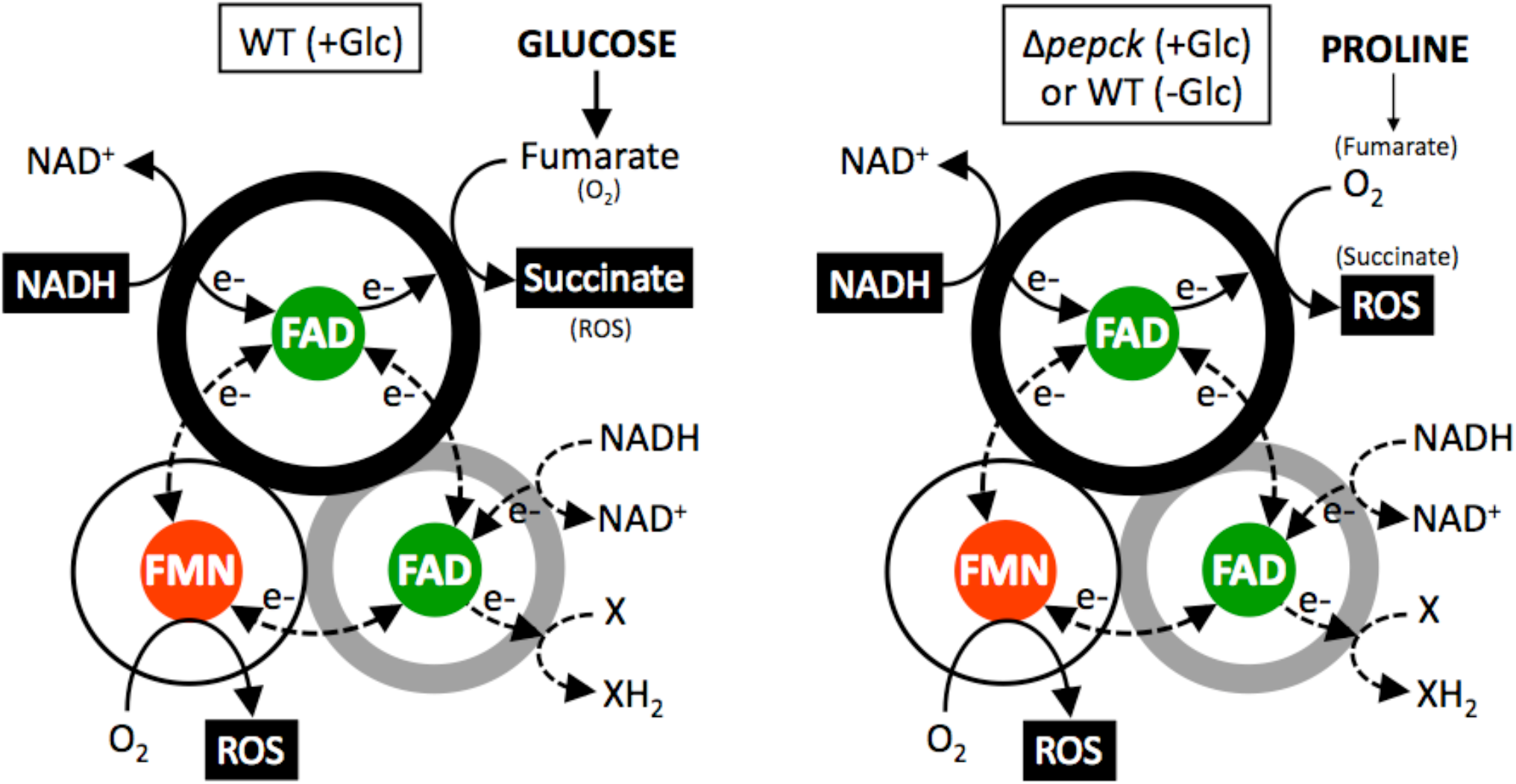
These schematic representations illustrate our hypothesis supported by our data and the previously described ability of FRDg to produce ROS(38). We propose that fumarate and oxygen compete for electrons provided by FAD in the FRD domain. The covalently bound flavin in the ApbE domain could also transfer electrons to oxygen. Unknown electron flows within FRDg and with potential substrates are indicated by dotted lines. The covalently bound flavin in the ApbE domain (FMN) and the FAD prosthetic cofactor in the FRD and Cytb5R domains (FAD) are shown in red and green respectively.

The next obvious question to ask is what would be the possible role of ROS production by the glycosomal fumarate reductase? ROS have historically been viewed as toxic metabolic by-products and causal agents of many pathologies. This notion is indeed supported by irreversible damages to cellular components caused by high levels of cellular ROS (39). However, this line of thinking has gradually shifted towards a more positive view with the growing body of evidence showing that lower levels of ROS are essential signals dictating biological outcomes, such as proliferation, adaptation and differentiation (40). Interestingly, Dolezelova *et al*. recently showed that cytosolic ROS trigger the differentiation of *T. brucei* procyclics into epimastigotes in the *in vitro* differentiation model based on overexpression of RNA-binding protein 6 (RBP6) (41, 42). Indeed, expression of a cytosolic catalase abolished differentiation of the parasite into mature epimastigotes upon induction of RBP6 expression, which was interpreted as the consequence of the degradation of hydrogen peroxide produced by the respiratory chain. Due to its membrane-permeability and high stability (43), hydrogen peroxide could be produced in any cell compartment, such as glycosomes, to exert its cytosolic effect. Therefore, FRDg could participate in the production of ROS used to trigger differentiation. This hypothesis is particularly relevant under the glucose-depleted conditions encountered in the midgut of the fly, which resembles the situation faced by the Δ*pepck* mutant grown in rich medium (see Fig S1), *i.e.* no contribution of FRDg in succinate production. We therefore propose that, in the glucose-depleted environment of the midgut of the fly, FRDg is mainly used for ROS production to participate in the differentiation of procyclics into epimastigotes. Our model is strengthened by the previous observation that the absence of FRDg (^*RNAi*^FRDg cell lines) does not affect growth of the PCF trypanosome in glucose-rich and glucose-depleted conditions, since the parasite has developed alternative means to maintain the glycosomal redox balance and excrete fumarate instead of succinate from glucose metabolism (14, 20, 21). This indeed suggests that the main role of FRDg is possibly not to participate in the glycosomal redox balance.

## Methods

### Trypanosomes, cell cultures and preparation of glycosomal fractions

The PCF of *T. brucei* EATRO1125.T7T (TetR-HYG T7RNAPOL-NEO) were cultivated at 27°C in the presence of 5% CO_2_ in SDM79 medium containing 10% (v/v) heat-inactivated fetal calf serum and 3.5 mg ml^-1^ hemin (44). Subcellular fraction enriched in glycosomes was prepared by differential centrifugation of parental and Δ*ppdk*/Δ*pepck*/^*RNAi*^GPDH.i PCF trypanosomes as described in (19), after homogenizing the cells with silicon carbide as grinding material. Briefly, 5 × 10^9^ cells were washed once in 50 ml of STE (25 mM Tris, 1 mM EDTA, 250 mM sucrose pH 7.8). After centrifugation, the pellet was resuspended in 0.5 ml of homogenization buffer STE (STE supplemented with ‘Complete EDTA-Free’ protease-inhibitor cocktail, Roche Applied Science, Mannheim, Germany) and ground in a pre-chilled mortar with 1.5 gr of wet-weight silicon carbide per gram of cell pellet. The cells were microscopically checked for at least 90% disruption. The cell lysate was diluted in 7 ml of homogenization buffer, centrifuged at 1000 g and then at 5000 g for 10 min each, at 4°C. The supernatant was centrifuged at 33,000 g for 10 min at 4°C to yield the glycosome-enriched pellet, which was resuspended in 2 ml of STE buffer and loading on a continuous sucrose gradient (1-2 M sucrose in STE). After centrifugation at 39,000 rpm in a vertical rotor, the band corresponding to glycosomes was collected, five-time diluted in STE and centrifuged at 33,000 g for 30 min at 4°C to yield the glycosomal pellet, which was resuspended in 0.5 ml of STE.

### Mutant cell lines

The single Δ*pepck::BSD*/Δ*pepck::PAC* (Δ*pepck*) and double Δ*ppdk*::TetR-HYG/Δ*ppdk*::T7RNAPOL-NEO Δ*pepck*::BSD/Δ*pepck::PAC* (Δ*ppdk*/Δ*pepck*) null mutant cell lines have been generated before (14, 18, 20, 24). RNAi-mediated inhibition of gene expression of the glycosomal glycerol-3-phosphate dehydrogenase gene (*GPDH*, EC 1.1.1.8, Tb927.8.3530) was performed in the Δ*ppdk*/Δ*pepck* PCF by expression of stem-loop “sense-antisense” RNA molecules of the targeted sequences corresponding to a 564 bp fragment (position 223 to 786) of the *GPDH* gene (45, 46), using the pLew100 expression vector, which contains the phleomycin resistance gene (kindly provided by E.Wirtz and G.Cross) (47). Similarly, the same approach was used to down-regulate expression of the chimeric FRDg-m2 isoform in the Δ*pepck\** cell line, by targeting a 544 bp fragment (position 1937 to 2480) of the *FRDm2* gene (Tb927.5.940). The resulting pLew100-GPDH-SAS and pLew100-FRDm2-SAS plasmids containing a sense and antisense version of the targeted gene fragment, separated by a 58-bp and 50-bp fragment, respectively, under the control of a PARP promoter linked to a prokaryotic tetracycline operator, were constructed as previously described using the HindIII, XhoI and BamHI restriction sites (18, 20). To express the FRDg isoform in the Δ*pepck* background, the *FRDg* gene (Tb927.5.930) was inserted in the HindIII and BamHI restriction sites of the pLew100 vector to produce pLew100-FRDg plasmid. The *FRDg-*Δ*SKI* recombinant gene coding for a FRDg isoform lacking the three C-terminal residues forming the PTS1 (SKI tripeptide) was generated by replacing the 233-bp ApaI/BamHI fragment of the pLew100-FRDg plasmid by the same fragment missing the 9 residues coding for the SKI tripeptide. The *FRDg-Δctl* recombinant gene coding for a FRDg isoform lacking the central FRD domain was generated by removing the 1361-bp PvuII/PspOMI fragment of the pLew100-FRDg plasmid, corresponding to position 997 bp and 2358 bp in the *FRDg* gene, followed by recircularization of the resulting plasmid. To produce the *FRDg-ΔNterm* recombinant genes a 2343-bp PCR fragment corresponding to position 1083 bp to 3426 bp of the *FRDg* gene was inserted into the HindIII and BamHI restriction sites of the pLew100 vector. To generate the *FRDg-ΔNterm/ctl* and *FRDg-Δ2-9* constructs, the 1504-bp HindIII/XhoI fragment of the pLew100-FRDg plasmid, encoding the first 498 amino acids of FRDg, was replaced by a 498-bp HindIII/XhoI fragment and a 1477-bp HindIII/XhoI fragment, respectively.

To constitutively express EGFP in the Δ*pepck\** cell line, the EGFP sequence was inserted in the HindIII and BamHI restriction sites of the pLew100 vector, which was modified by removing the two tetracycline operator sequences. The pLew100-FRDg-m2-SAS, pLew100-FRDg and pLew100-EGFPct plasmids designed to generate the Δ*pepck\**/^*RNAi*^FRDg-m2, Δ*pepck\**/^*OE*^FRDg and Δ*pepck\**/^*OE*^EGFPct cell lines were provided by the Genecust company. The plew100 recombinant plasmids were linearized with the restriction enzyme NotI and transfected into the Δ*ppdk*/Δ*pepck* (pLew100-GPDH-SAS), Δ*pepck\** (all the other plasmids) or parental (pLew100-FRDg-ΔSKI) cell lines.

Selection of all these mutant cell lines was performed in SDM79 medium containing hygromycin (25 μg ml^-1^), neomycin (10 μg ml^-1^), blasticidin (10 μg ml^-1^), puromycin (1 μg ml^-1^) and/or phleomycin (5 μg ml^-1^). Aliquots were frozen in liquid nitrogen to provide stocks of each line that had not been in long-term culture. Induction of RNAi cell lines was performed by addition of 1 μg ml^-1^ tetracycline.

### Competitive growth assay

The objective of this assay is to determine slight but significant doubling time difference between a conditional mutant and an EGFP tagged reference cell line, upon co-culture experiments. This assay is based on the co-culture of a tetracycline-inducible conditional mutant cell line and a reference cell line constitutively expressing EGFP (Δ*pepck\**/^*OE*^EGFPct_F5), which has a doubling time of 14.26 ±1.06 h. The SDM79 medium was inoculated with the Δ*pepck\**/^*OE*^EGFPct_F5 reference cell line (1.4 × 10^6^ cells ml^-1^) and a mutant cell line (0.6 × 10^6^ cells ml^-1^), in the presence or the absence of 1 mg ml^-1^ tetracycline, and the proportion of EGFP-positive cells was determined every day by flow cytometry using a Guava EasyCyte Flow Cytometer (Merck Millipore). The difference of the percentage of EGFP negative cells between induced and non-induced conditions were plotted as a function of time of growth, in order to estimate the growth difference between the non-induced and tetracycline-induced cell line.

### Western blot analyses

Total protein extracts (3-5 × 10^6^ cells) or glycosomal extracts of the parental (EATRO1125.T7T) or mutant PCF of *T. brucei* were separated by SDS-PAGE (8% or 10%) and immunoblotted on TransBlot Turbo Midi-size PVFD Membranes (Bio-Rad) (48). Immunodetection was performed as described (48, 49) using as primary antibodies the rabbit anti-FRD (aFRD, 1:100) (21), the rabbit anti-FRDg (aFRDg, 1:100) (22), the rabbit anti-FRDm2 (aFRDm2, 1:100, produced by Proteogenix from the EISKSVFPDASLGV and ELGHNKSNIVTL peptides), the rabbit anti-PEPCK (aPEPCK, 1:1000) (20), the rabbit anti-GPDH (aGPDH, 1:1000) (50), the rabbit anti-PPDK (aPPDK, 1:1000) (51), the rabbit anti-enolase (aENO 1:100,000, gift from P. Michels, Edinburgh, UK), the rabbit anti-GAPDH (aGAPDH 1:10,000, gift from P. Michels, Edinburgh, UK), the rabbit anti-PFK (aPFK 1:5,000, gift from P. Michels, Edinburgh, UK) and the rabbit antibody against the glycosomal isocitrate dehydrogenase, anti-IDHg (aIDHg 1:20,000, produced by Pineda (Berlin, Germany) against recombinantly expressed full-length IDHg). Anti-rabbit IgG conjugated to horseradish peroxidase (Bio-Rad, 1:5,000 dilution) was used as secondary antibody. Detection was performed using the Clarity Western ECL Substrate as described by the manufacturer (Bio-Rad). Images were acquired and analyzed with the ImageQuant LAS 4,000 luminescent image analyzer. For near-infrared fluorescent Western blotting the mouse anti-PFR-A/C (aPFR, 1:2,000) (52) and the rabbit anti-FRD (aFRD, 1:1,000) (21) were used as primary antibodies, the IR-BLOT 800 anti-mouse IgG (Cyanagen Srl, 1:5,000 dilution) and IRDye 680LT anti-rabbit IgG (LI-COR Bioscience, 1:5,000 dilution) as secondary antibodies. Image acquisition was performed with the Odyssey CLx Near-Infrared Fluorescence Imaging System and the dedicated software Image Studio (LI-COR Bioscience).

### Analysis of FRDg flavinylation

As described in (53), gels resulting from SDS-PAGE were scanned with a Typhoon TRIO Variable Mode Imager System (GE Healthcare) at λex = 488 nm and λem = 526 nm for detection of covalently bound flavin and at λex = 670 nm and λem = 633 nm for visualization of the Blue Prestained Protein Standard (NEB).

### Digitonin permeabilization

Digitonin permeabilization was performed as described before (18). Briefly, trypanosomes were washed two times in cold PBS and resuspended at 6.5 ′ 10^8^ cells ml^-1^ (corresponding to 3.3 mg of protein per ml) in STE buffer (250 mM sucrose, 25 mM Tris, pH 7.4, 1 mM EDTA) supplemented with 150 mM NaCl and the Completeä Mini EDTA-free protease inhibitor cocktail (Roche Applied Bioscience). Cell aliquots (200 ml) were incubated with increasing quantities of digitonin (Sigma) for 4 min at 25°C, before centrifugation at 14,000 *g* for 2 min to collect the cellular pellet.

### Enzymatic activities

Sonicated (5 sec at 4°C) crude extracts of trypanosomes resuspended in cold hypotonic buffer (10 mM potassium phosphate, pH 7.8) were tested for enzymatic activities. NADH-dependent FRD, glycerol kinase and malic enzyme activities were measured at 340 nm via oxidation of NADH or NADPH, according to published procedures (18).

### Southern blot analysis

Genomic DNA (10 μg) from the parental (EATRO1125.T7T) and Δ*ppdk*/Δ*pepck*/^*RNAi*^GPDH cell lines, extracted as previously described (54), was digested with the NcoI, PvuI, NdeI or XhoI restriction enzymes, separated by electrophoresis in a 0.8% agarose gel and transferred onto a nylon membrane (Hybond N^+^, Roche Molecular Biochemicals). The membrane was hybridized with digoxigenin-labeled DNA probes synthesized with a PCR DIG probe synthesis kit (Roche Molecular Biochemicals) as recommended by the supplier. The *FRD* probe was generated by PCR amplification, using the primer pair 5’- GTGTAACGTCGTTGCTCAGTGAGA-3’ / 5’- GCGAAATTAAATGGGCCCCGCGACG-3’. Probe-target hybrids were visualized by a chemiluminescent assay with the DIG luminescent detection kit (Roche Molecular Biochemicals), according to the manufacturer’s instructions. Blots were exposed to ImageQuant LAS4010 (GE Healthcare Life Sciences) for approximately 20 min.

### Label-free quantitative proteomics

Total extracts and glycosome-enriched fractions of trypanosomes were loaded on a 10% acrylamide SDS-PAGE gel and proteins were visualized by Colloidal Blue staining. For total extracts, migration was performed classically and each protein lane was cut into 4 equal segments. For the glycosome-enriched fractions, migration was stopped when samples had just entered the resolving gel and the unresolved region of the gel was cut into only one segment. Finally, each SDS-PAGE band was cut into 1 mm × 1 mm gel pieces. Protein digestion and nano-liquid chromatography–tandem mass spectrometry analyses on LTQ Orbitrap XL were performed as previously described (16). For protein identification, Sequest HT and Mascot 2.4 algorithms through Proteome Discoverer 1.4 Software (Thermo Fisher Scientific Inc.) were used for protein identification in batch mode by searching against a *Trypanosoma brucei* protein database (11 119 entries, release 46). This database was downloaded from http://tritrypdb.orgwebsite. Two missed enzyme cleavages were allowed. Mass tolerances in MS and MS/MS were set to 10 ppm and 0.6 Da. Oxidation of methionine, acetylation of lysine and deamidation of asparagine and glutamine were searched as dynamic modifications. Carbamidomethylation on cysteine was searched as static modification. Peptide validation was performed using Percolator algorithm (55) and only “high confidence” peptides were retained corresponding to a 1% False Discovery Rate (FDR) at peptide level. Raw LC-MS/MS data were imported in Progenesis QI (version 2.0; Nonlinear Dynamics, a Waters Company) for feature detection, alignment, and quantification. All sample features were aligned according to retention times by manually inserting up to fifty landmarks followed by automatic alignment to maximally overlay all the two-dimensional (m/z and retention time) feature maps. Singly charged ions and ions with higher charge states than six were excluded from analysis. All remaining features were used to calculate a normalization factor for each sample that corrects for experimental variation.

Peptide identifications (with FDR<1%) were imported into Progenesis. Only non-conflicting features and unique peptides were considered for calculation of quantification at protein level, a fold changes above 2 The mass spectrometry proteomics data have been deposited to the ProteomeXchange Consortium via the PRIDE (56) partner repository with the dataset identifier PXD020185.

### Analysis of excreted end products from the metabolism of glucose and proline by proton NMR

2 × 10^7^ *T. brucei* PCF were collected by centrifugation at 1,400 g for 10 min, washed once with phosphate-buffered saline (PBS) and incubated in 1 ml (single point analysis) of PBS supplemented with 2 g l^-1^ NaHCO_3_ (pH 7.4). Cells were maintained for 6 h at 27°C in incubation buffer containing 4 mM [U-^13^C]-glucose and 4 mM non-enriched proline. The integrity of the cells during the incubation was checked by microscopic observation. The supernatant (1 ml) was collected and 50 μl of maleate solution in D_2_O (10 mM) was added as internal reference. H-NMR spectra were performed at 500.19 MHz on a Bruker Avance III 500 HD spectrometer equipped with a 5 mm cryoprobe Prodigy. Measurements were recorded at 25°. Acquisition conditions were as follows: 90° flip angle, 5,000 Hz spectral width, 32 K memory size, and 9.3 sec total recycle time. Measurements were performed with 64 scans for a total time close to 10 min 30 sec. Resonances of the obtained spectra were integrated and metabolites concentrations were calculated using the ERETIC2 NMR quantification Bruker program.

## Acknowledgements

We thank Paul A. Michels (Edinburgh, Scotland) for providing us with the anti-GAPDH, anti-PFK and anti-enolase immune sera and Marc Ouellette for critical reading of the manuscript.

## Funding and additional information

This work was supported by the Centre National de la Recherche Scientifique (CNRS), the Université de Bordeaux, the ParaMet PhD programme of Marie Curie Initial Training Network, the Agence Nationale de la Recherche (ANR) through GLYCONOV and ADIPOTRYP grants of the “Générique” call, the Laboratoire d’Excellence (LabEx) ParaFrap ANR-11-LABX-0024. Work in the Munich lab was supported by a student research fellowship and the BioNa junior career award of the Faculty of Biology of LMU to R.S. and S.B., respectively.

---

**Fig S1.**
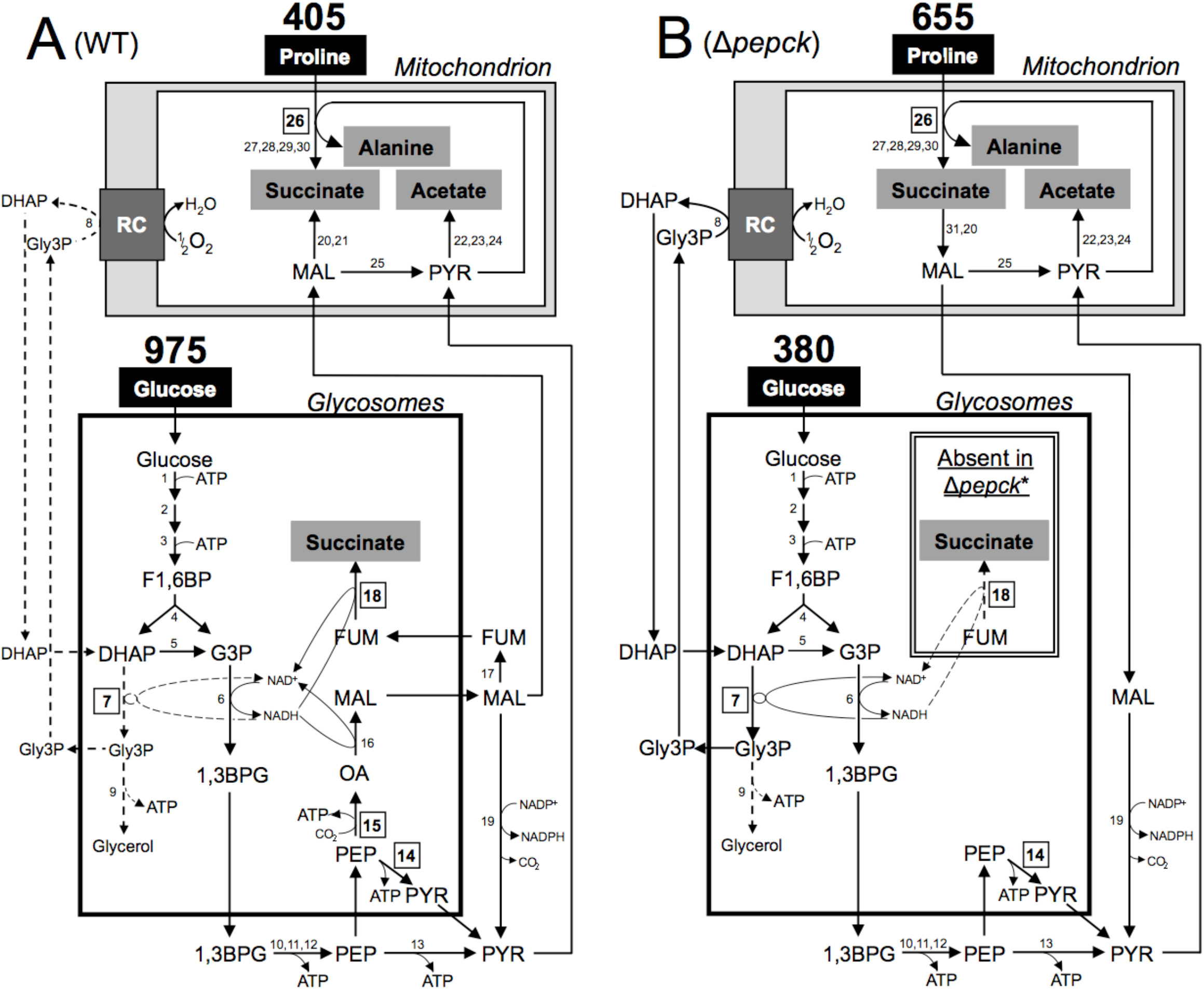
Schematic representation of the central metabolism of the wild-type and Δ*pepck* mutant PCF cell lines in glucose-rich medium. This figure highlights the catalytic steps from glucose and proline metabolism in the wild-type (WT) (panel A), Δ*pepck* and Δ*pepck*\* (panel B) cell lines. The mitochondrial pathways have been considerably simplified, in particular the respiratory chain (RC) represented by a grey box. Excreted end products from degradation of glucose and proline are in a grey background and dashed lines represent enzymatic steps not used or used at a background noise level. Boxed numbers correspond to the enzymatic steps investigated. The rates of glucose and proline consumption (nmol h^−1^ mg^−1^ of protein) indicated above the carbon i names are deduced from Tables S1 and are consistent with previous data (18). The double box inside the glycosomes including the FRDg step is missing in the Δ*pepck*\* cell line. Abbreviations: 1,3BPG, 1,3-biphosphoglycerate; DHAP, dihydroxyacetone phosphate; F1,6BP, fructose 1,6-bisphosphate; FUM, fumarate; G3P, glyceraldehyde 3-phosphate; Gly3P, glycerol 3-phosphate; MAL, malate; OA, oxaloacetate; PEP, phosphoenolpyruvate; PYR, pyruvate; RC, respiratory chain. Enzymes: 1, hexokinase; 2, glucose-6-phosphate isomerase; 3, phosphofructokinase; 4, aldolase; 5, triose-phosphate isomerase; 6, glyceraldehyde-3-phosphate dehydrogenase; 7, glycosomal NADH-dependent glycerol-3-phosphate dehydrogenase (GPDH); 8, mitochondrial FAD-dependent glycerol-3-phosphate dehydrogenase (GPDH); 9, glycerol kinase; 10, phosphoglycerate kinase; 11, phosphoglycerate mutase; 12, enolase; 13, pyruvate kinase; 14, pyruvate phosphate dikinase (PPDK); 15, phosphoenolpyruvate carboxykinase (PEPCK); 16, glycosomal malate dehydrogenase; 17, fumarase; 18, glycosomal NADH-dependent fumarate reductase (FRDg); 19, cytosolic malic enzyme; 20, mitochondrial malate dehydrogenase; 21, mitochondrial NADH-dependent fumarate reductase (FRDm1); 22, pyruvate dehydrogenase complex; 23, acetate:succinate CoA-transferase; 24, acetyl-CoA thioesterase; 25, mitochondrial malic enzyme; 26, proline dehydrogenase (PRODH); 27, pyrroline-5 carboxylate dehydrogenase; 28, alanine aminotransferase; 29, α-ketoglutarate dehydrogenase complex; 30, succinyl-CoA synthetase; 31, succinate dehydrogenase.

**Fig S2.**
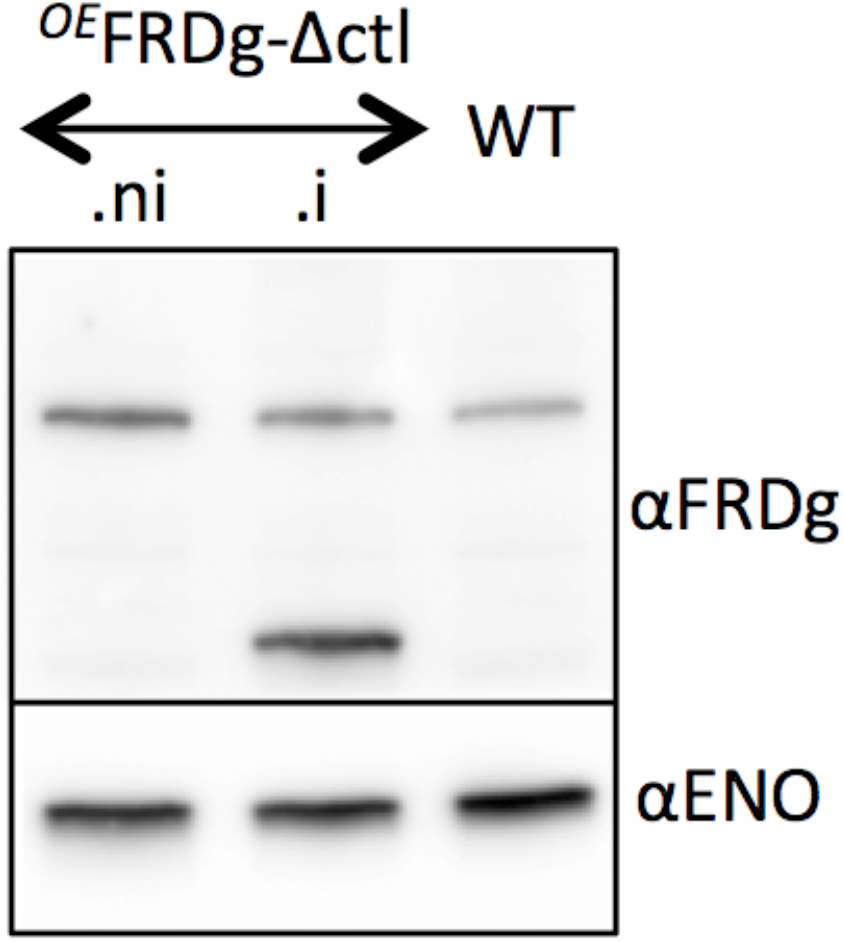
Expression of FRDg-Δctl in the parental (WT) background.

